# Hybrids reveal accessible chromatin *trans* genetic associations

**DOI:** 10.1101/2025.10.09.681407

**Authors:** Mark A.A. Minow, Alexandre P. Marand, Xuan Zhang, Cullan A. Meyer, Jordan Pelkmans, Robert J. Schmitz

**Affiliations:** Department of Genetics, University of Georgia; 120 E Green St, Athens, GA 30602, USA; Department of Molecular, Cellular, and Developmental Biology, University of Michigan; 1105 N University Ave, Ann Arbor, MI, USA; Nell Hodgson Woodruff School of Nursing, Emory University; 1520 Clifton Rd NE, Atlanta, GA 30322, USA

## Abstract

Crosses between genetically distant *Zea mays* (Maize) inbreds cause hybrid vigour (heterosis): an over-dominant increase in vegetative growth and grain yield. Although the contributing loci are unknown, heterosis strength varies polygenically, with crosses between stiff stalk and non-stiff stalk lineages often producing the highly heterotic F1 seed used for agriculture. As F1 genomes conserve the parental *cis*-regulatory state, F1 *trans*-interactions between haplotypes underpin F1 trait over-dominance and heterotic gene expression. Gene expression relies upon transcription factors binding *cis*-regulatory elements in accessible chromatin regions (ACRs) and recruiting RNA POLYMERASE II. Here, assay for transposase accessible chromatin sequencing (ATAC-seq) examined *cis*-regulatory element chromatin accessibility inheritance in three different F1 parent-offspring pairs, including a representative stiff stalk-by-non-stiff stalk (B73xOh43) cross. Using concatenated reference alignment, we categorized ACR modes of inheritance (additive, dominant, under/over-dominant), revealing that ACRs are largely inherited additively, and, concordantly, that chromatin accessibility has strong narrow-sense heritability. However, mirroring other heterotic phenotypes, the B73xOh43 cross exhibited the most dominant and over-dominant inheritance. Comparing allele-specific parent-to-offspring chromatin accessibility, we identified F1 differentially *trans*-regulated ACRs. Using a matched chromatin accessible diversity panel, we found B73xOh43 SNPs with more ACR *trans*-associations than expected due to chance. These *trans*-regulatory variants revealed how cell-division and anabolism are elevated in heterotic F1 seedlings. This multiple discovery filtering (MDF) approach to *trans*-regulatory variant detection is genomic modality agnostic and stands to empower *trans*-relationship dissection in diverse genetic models.

## Introduction

Both differing environments and individual genetic diversity influence trait variation, with the heritability of a phenotype representing the portion of genetic control. Many highly heritable and important traits are influenced by the inheritance of many unlinked loci. For example, in *Zea mays* (maize) these complex, quantitative or polygenic traits include plant architecture(Wallace *et al*., 2014; Wang *et al*., 2023*a*), flowering time(Buckler *et al*., 2009; Li *et al*., 2016; Romero Navarro *et al*., 2017) and yield(Mural *et al*., 2022). Since quantitative traits encompass such important agronomic phenotypes, there is much interest in understanding the underlying genetics. However, the complexity of the gene regulatory networks underpinning quantitative traits obfuscates the mechanistic dissection of the pathways controlling traits.

In contrast, the inheritance of functionally distinct alleles is well understood. Most alleles equally control their respective phenotypes, resulting in additive inheritance. For example, under controlled environments, if two parents are homozygous for two different additive alleles (AA x aa), the progeny (Aa) would have a phenotype resembling their average, or mid-parental value. However, non-additive modes of inheritance also occur. If the progeny phenotype mirrors that of one parent, it is known as dominant inheritance. Alternatively, progeny can inherit a transgressive phenotype, higher or lower than either parent, resulting from over-dominance or under-dominance respectively.

From mules to maize, diverse eukaryotes exhibit an interesting form of over-dominance: heterosis (hybrid vigour) (Kaeppler, 2012). Heterosis follows a ‘wide’ cross between two distantly related individuals, causing simultaneous over-dominance for many phenologically disparate phenotypes. However, hybrid vigour strength is genetically variable, making heterosis itself a variable quantitative trait (Stuber *et al*., 1992; Kaeppler, 2012; Riedelsheimer *et al*., 2012; Li *et al*., 2018; Xiao *et al*., 2021; Wang *et al*., 2023*b*). Strong and uniform filial one (F1) heterosis is exploited in maize agronomy (Shull, 1908; East and Jones, 1920; Duvick, 2005), often involving crosses between inbreds from different heterotic groups – notably crosses between stiff stalk and non-stiff stalk lineages. These F1 plants exhibit over-dominance for many traits (Xiao *et al*., 2021), including vegetative seedling growth (Li *et al*., 2018), which relates to yield (Lu *et al*., 2011; Trachsel *et al*., 2017).

Vegetative growth requires water, minerals, carbon skeletons and energy – with the latter two components being supplied by photosynthesis. Photosynthates (sugars) are produced in ‘source’ chlorophyllic tissues and then translocated to consumptive ‘sink’ tissues, like organs undergoing growth (MacNeill *et al*., 2017). Intuitively, elevated photosynthesis increases growth (South *et al*., 2019; Wu *et al*., 2019). However, increased sink strength can also improve crop growth (Nuccio *et al*., 2015)(Nuccio et al., 2015), partly by increasing carbon export from source tissues, alleviating the negative photosynthetic feedback driven by high sugar concentrations (Moore *et al*., 2003; Cho *et al*., 2006; Vanderwall and Gendron, 2023). During vegetive growth, leaf production is a major carbohydrate sink that doubles as an energetic investment – a current carbon cost yielding future photosynthetic capacity (Wright *et al*., 2004). Carbon investments in organ formation follow a pattern of inter-trait correlations known as the acquisitive–conservative axis, which varies between (Wright *et al*., 2004) and within (Gorné *et al*., 2022) species and relates to organ dry mass per volume. Conservative tissues are high cost, but long lived and resilient, providing photosynthetic extended ‘returns.’ In contrast, acquisitive tissues are low cost, more aggressively using water, carbon and minerals to increase compounding photosynthetic returns, but trade-off herbivory/disease/stress susceptibility, reducing organ lifespans. Crop growth may be too conservative, as the negatives associated with acquisitive growth are ameliorated by agronomic practices (e.g. pesticides), transgenesis (e.g. *Bt*-toxin(Santos-Amaya *et al*., 2015)), and financial instruments (crop insurance). However, the molecular mechanisms promoting acquisitive vegetative vigour are complex and poorly understood, hindering engineering approaches to improve crop growth and yield (Simmons *et al*., 2021). Likewise, interspecific comparisons suggest conventional breeding has not elevated crop acquisitional traits beyond that found in pre-domesticate relatives (Gómez-Fernández *et al*., 2024). Studying heterotic vegetative vigour may help reveal the genetic underpinnings of growth rates more generally. Although common heterotic regulators, like those around the circadian clock (Ni *et al*., 2009; Yang *et al*., 2021) and floral regulator genes (Xiao *et al*., 2021), are emerging, the polygenic and phenological nature of hybrid vigour leaves its mechanisms mysterious.

Heterosis is thought to involve altered transcription, indeed known heterotic strength determinants are transcription factors (TFs) (Ni *et al*., 2009; Xiao *et al*., 2021; Yang *et al*., 2021). TFs regulate gene transcription by binding specific DNA sequence motifs, or *cis*-regulatory elements (CREs), linked to genes(Schmitz *et al*., 2021; Marand *et al*., 2023). Bound TFs regulate RNA POLYMERASE II recruitment, coordinating transcription with developmental and environmental cues. TF binding mostly requires direct peptide-to-DNA interaction which is sterically restricted to nucleosome free (accessible) chromatin; these accessible chromatin regions (ACRs) can be uncovered via Assay for Transposase-Accessible Chromatin sequencing (ATAC-seq) (Buenrostro *et al*., 2013; Minnoye *et al*., 2021), revealing the CRE complement available for TF binding. Beyond variable CRE accessibility, TF activity is influenced by cellular biochemistry, with protein-protein interactions(Kim and Wysocka, 2023), hormone/metabolite binding (Carthew, 2021; Hornisch and Piazza, 2025) and post-translational modification(Weidemüller *et al*., 2021). Dissecting heterotic gene expression is challenging, with divergent CRE variants and protein complements acting in unison.

However, inbred-derived F1 hybrids are uniquely poised to empower the study of gene regulation, as the inbred haplotypes remain completely intact. Beyond allowing easy and exact genotypic replication, unrecombined haplotypes preserve the *cis*-regulatory environment that existed in the parental state; that is, global linkage between CRE variants and their target genes remains unchanged. In contrast, *trans*-regulation, defined as all regulatory factors unlinked to their gene targets, changes in the hybrid. TFs and other regulatory proteins from both parental states influence the F1 genome, with the *trans*-regulatory space encompassing that of both inbreds, albeit at a halved dose. As all *cis* effects are fixed, the mechanisms behind heterosis must lie in the F1 *trans*-regulatory differences; i.e. inter-chromosomal genetic interactions ultimately manifest heterotic phenotypes. However, delineation of these *trans*-regulatory relationships between genes remains technically challenging. Intra-specific *trans*-regulatory variation has small expression effect sizes (Dixon *et al*., 2007; Göring *et al*., 2007; West *et al*., 2007; Swanson-Wagner *et al*., 2009; Li *et al*., 2013; Võsa *et al*., 2021; Zhan *et al*., 2025), which, when combined with an extremely high multiple testing burden, precludes high-confidence association between genetic variants and their unlinked targets.

Transcriptional decisions are critical ‘integration points’ in diverse processes (Katagiri and Chua, 1992; Paz-Ares *et al*., 2002; Lambert *et al*., 2018) and much effort has been placed into the study of what controls transcription, both at a single gene level and globally. Heritability estimates, both narrow-sense (h^2^; assuming only additive genetic effects)(Monks *et al*., 2004; Dixon *et al*., 2007; Göring *et al*., 2007; Price *et al*., 2011; Yang *et al*., 2014; Ouwens *et al*., 2020) and broad-sense (H^2^; encompassing all genetic effects, including dominance and epistasis) (Sun *et al*., 2023), indicate that transcription is under genetic control. In contrast to transcription, few measures of chromatin accessibility heritability exist(Gate *et al*., 2018; Zhu *et al*., 2024), and questions remain surrounding how much the environment or genetics affect nucleosome occupancy. However, measurements of chromatin accessibility within genetically diverse mapping populations have revealed chromatin accessibility quantitative trait loci (caQTL) (Gate *et al*., 2018; Kumasaka *et al*., 2019; Aygün *et al*., 2023; Benaglio *et al*., 2023; Zeng *et al*., 2024; Marand *et al*., 2025; Zhu *et al*., 2025), highlighting that genetic variation controls the accessibility of certain CREs. Yet, due to the aforementioned difficulties in mapping *trans*-regulatory differences, known caQTL mainly encompass *cis* (linked) genetic determinants (Gate *et al*., 2018; Kumasaka *et al*., 2019; Aygün *et al*., 2023; Zeng *et al*., 2024; Marand *et al*., 2025).

Here, concurrently grown seedlings from a trio of inbreds, B73, Oh43 and Ki3, and their corresponding F1s, B73 x Ki3, B73 x Oh43, Oh43 x Ki3 (hereafter BxK, BxO, OxK), were subjected to chromatin accessibility profiling via bulk ATAC-seq. Using the corresponding reference genomes(Hufford *et al*., 2021), we comprehensively measured haplotype-resolved ACR complements, describing intra-specific ACR conservation and divergence. Comparison of parental and F1 offspring chromatin accessibility revealed the modes and prevalence of ACR inheritance and enabled chromatin accessibility h^2^ estimates. Exploiting F1 *cis*-effect preservation and allele-specific ATAC-reads, we discovered loci with accessibility altered by the F1 *trans*-regulatory environment. To investigate the genetic underpinnings of this F1 ACR *trans*-regulation, we combined these bulk measurements with a tissue and environmentally matched single-cell chromatin accessibility diversity panel(Marand *et al*., 2025). To empower *trans*-effect genetic mapping, we developed and showcase a novel Multiple Discovery Filtering (MDF) approach that controls for false positives (type I errors) by uncovering *trans*-regulatory variants with many unlinked ACR/gene associations. This approach discovered both *trans*-regulatory variants and biological processes associated with BxO seedling heterosis.

## Results

### Characterizing F1 Chromatin Accessibility Inheritance

Chromatin accessibility from seven-day old B73, Oh43, Ki3, BxK, BxO, and OxK seedlings were measured via bulk ATAC-seq (Figure 1A) with genotypic replication between four and six. This tissue was chosen to be environmentally and developmentally matched to a larger, single-cell chromatin accessibility diversity panel(Marand *et al*., 2025). The inbred genotypes were selected as they: 1) are well distributed on the maize phylogeny (Hu *et al*., 2021) 2) have reference genome assemblies(Hufford *et al*., 2021) 3) vary in heterosis, with the BxO F1 (stiff stalk x non-stiff stalk)(Li *et al*., 2018) having stronger heterosis than the other two F1s. To most accurately measure the differences in chromatin accessibilities between haplotypes, we used both a single (B73) reference (Figure 1B) and a concatenated genome alignment approach(Engelhorn *et al*., 2025) that enables detection of presence-absence variant ACRs and abrogates haplotype mapping bias. Quality control metrics supported good ATAC-seq quality (Table S1), but one B73 library was excluded due to low read number. For each concatenated reference, Tn5 insertion peaks, or ACRs, were identified if an overlapping peak was detected in n-1 biological replicates, producing BxK, BxO and OxK reference based ACRs, all with equivalent haplotype peak distribution (BxK: 51,459 B73 and 51,060 Ki3 ACRs; BxO: 40,939 B73 and 40,649 Oh43 ACRs; OxK: 63,845 Oh43 and 63,048 Ki3 ACRs)(Figure 1C; Table S2). Next, in each F1, these ACRs were categorized (Data S1,2,3) based on their proximity to genes (TSS, proximal if <2kb upstream of TSS, intergenic if >2kb upstream TSS or genic). All ACR sets had similar distributions of ACR-to-gene distance (Figure 1D and E). We also classified all ACRs based on their conservation status between haplotypes, with, 61.5-65% of ACRs being shared (high sequence identity) between haplotypes, 13.1-17.2% exhibiting ambiguous (poor) inter-haplotype sequence identity, 17.3-19.7% being unique to a haplotype and 2.2-2.8% duplicated, copy number variant (CNV) ACRs (Figure 1F). However, our CNV ACR detection is an underestimate, as limited multimapping (max 2 alignments) precludes the detection of evolutionarily young ACR large copy duplications containing high sequence identity.

**Figure 1.**
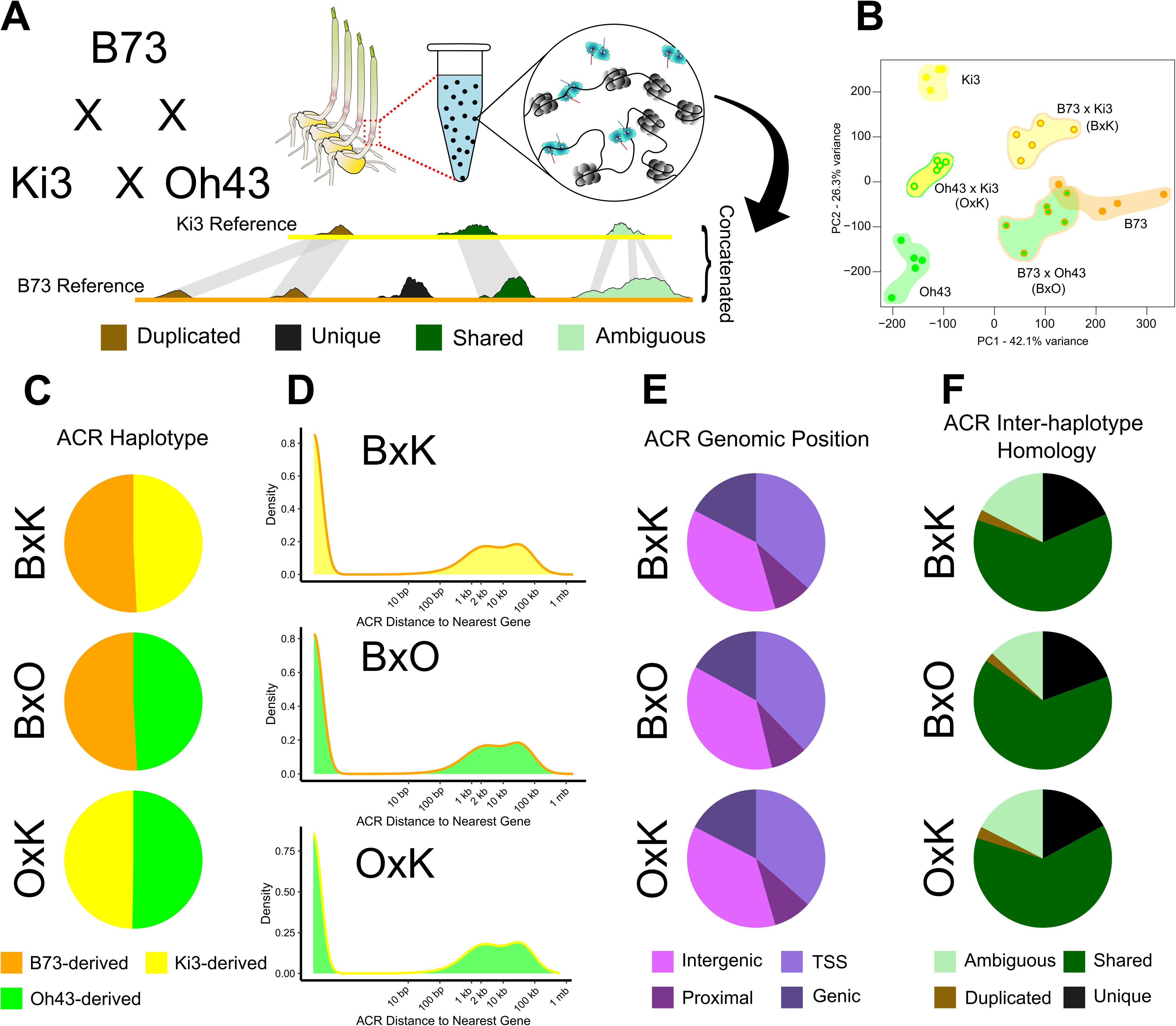
Characterizing parent-offspring chromatin accessibility via ATAC-seq with three concatenated reference alignments. (**A**) A trio of inbreds, B73, Oh43 and Ki3 were crossed to produce three F1s: B73 x Ki3 (BxK), B73 x Oh43 (BxO), and Oh43 x Ki3 (OxK). ATAC-seq chromatin accessibility measurements of the dissected first node (red dashed box) from seven-day old seedlings (n=4-6). These genotypes were grown under fixed environments to ascertain patterns of F1 chromatin accessibility inheritance. Data from the three parent-offspring pairs were aligned to a single reference (B73 – not shown) and concatenated reference genomes composed of the two parent haplotypes. Comparing accessible chromatin region (ACR) complements across haplotypes, we establish inter-haplotype homology groups: duplicated (brown), unique (black), shared (ACRs reciprocally exhibit 40% or more alignment between haplotypes) and ambiguous (ACRs unidirectionally exhibit 40% or more alignment between haplotypes). This revealed haplotype unique ACRs, duplicated ACRs, ACRs shared and conserved between haplotypes, and ambiguous ACRs, with poor inter-haplotype homology. (**B**) ATAC-seq data clustered by genotype, recapitulating the known F1 parent-offspring relationships. The ACR complement produced in the BxK, BxO and OxK concatenated references exhibited (**C**) balanced ACR haplotype representation, and (**D,E**) similar ACR distributions relative to genes. (**F**) Likewise, all genotypes exhibited similar representations of ACR inter-haplotype homology categories.

After defining the ACR complements on all haplotypes, we adapted a parent-offspring comparison(Coolon *et al*., 2014) to interrogate how chromatin accessibility (Tn5 insertion rates) varied from their parental states (Figure 2A-C; Figure S1), classifying ACR chromatin accessibility inheritance patterns as similar (invariable chromatin accessibility), additive, dominant, under-dominant, or over-dominant. Many ACRs (42.1-49.5%) were similarly accessible between parent and offspring, suggesting F1 conserved chromatin accessibility control over these loci. Of the ACRs exhibiting chromatin accessibility differences, additive inheritance was most frequent (38.5-49.9%), dominant inheritance patterns were abundant (6.8-11.0%), and over/under-dominance rare (0.2-0.8% and 0-0.02% respectively). Curiously, our parent-offspring accessibility plots differed in structure than those observed for gene expression(Coolon *et al*., 2014), with the ACR plot exhibiting long additive quadrant ‘tails’. We rationalized that this might be due to chromatin accessibility inheritance differing from mRNA inheritance. Specifically, we hypothesized that chromatin accessibility is more likely to exhibit simpler inheritance than gene expression, with more ACRs having binary, presence-absence variation driven entirely by *cis*-effects. In this case, the F1s would be expected to have half the chromatin accessibility of the parent, mirroring the halved allele dose. To examine this possibility, we filtered for ACRs with chromatin accessibility similar (one sample t-test, not FDR corrected p>0.05) to their half parental level, which recapitulates these additive quadrant ‘tails’ (Figure S2). Moreover, these ‘half-parental’ ACRs composed 19.0-38.1% of all additively inherited ACRs, indicating events of strictly allele-dose driven inheritance are common. Taken together, chromatin accessibility has inheritance like transcription, except with more prevalent high-fidelity allele-dose driven inheritance.

**Figure 2.**
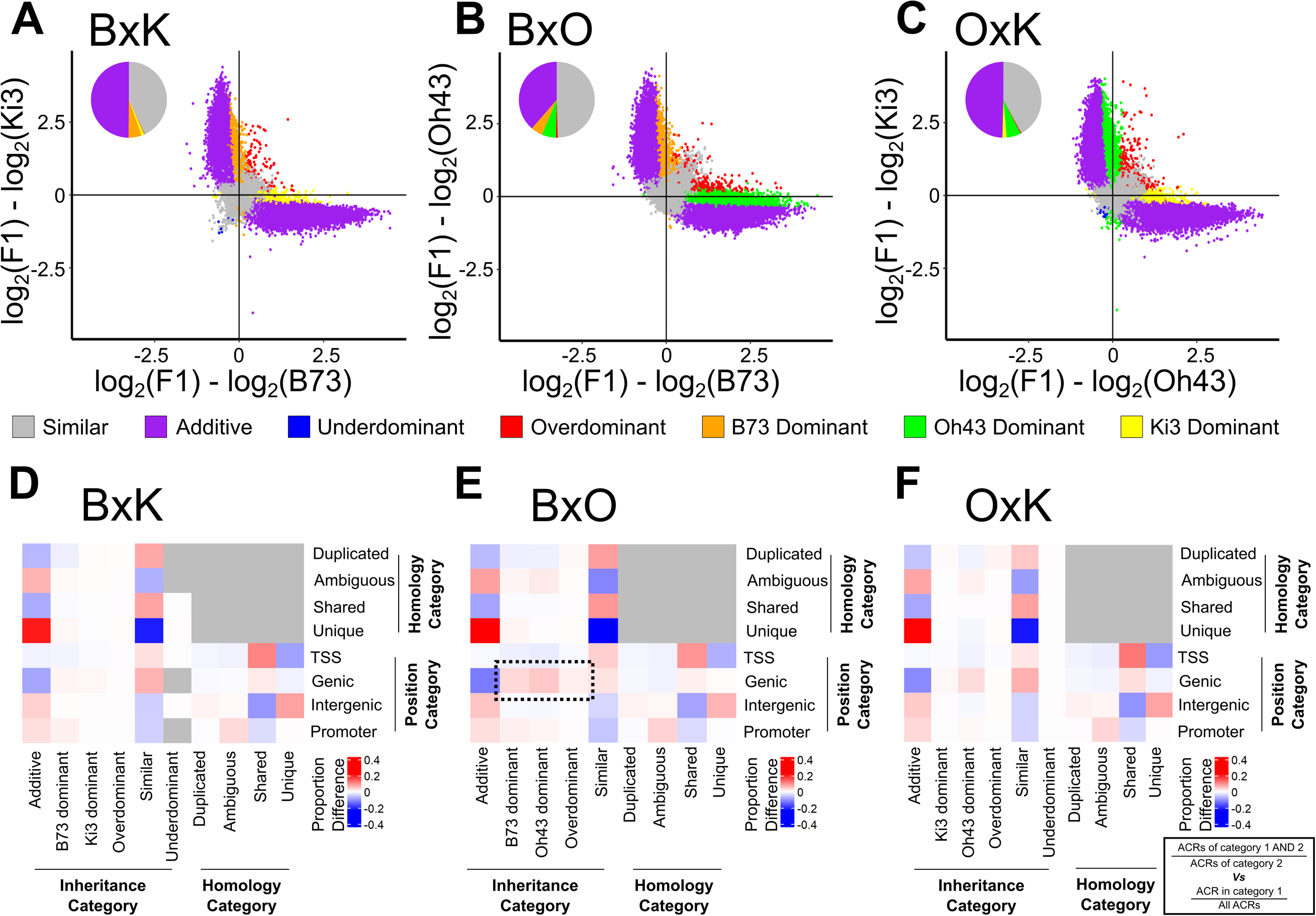
Modes of chromatin accessibility inheritance in three maize F1s. Parent-to-offspring comparison of chromatin accessibility revealed ACR mode of inheritance in (**A**) BxK, (**B**) BxO and (**C**) OxK. Of the ACRs exhibiting accessibility variation, most were additively inherited across all F1s. The most heterotic F1, BxO, exhibited the most dominant and over-dominant ACR inheritance and zero under-dominant ACRs. Proportional enrichment/depletion of ACR genomic position, inter-haplotype homology and mode of inheritance categories revealed similar patterns in (**D**) BxK, (**E**) BxO and (**F**) OxK. However, BxO (over)dominant ACRs had noticeably stronger enrichments (dashed box) over genic regions than observed in the other F1s. All enrichment/depletion analysis compared the proportion of overlap between two categories to the rate of category occurrence in the global ACR set (Solid black box). For example, the proportion of BxK Additive AND (∩) shared ACRs were compared to the proportion of additive ACRs in entire BxK ACR complement.

We also compared ACR inheritance patterns between the F1 sets. A similar number of dominant ACRs were observed on both B73 (BxK 5,639; BxO 4,088 ACRs) and Oh43 (BxO 4,922; OxK 7,831 ACRs) haplotypes, but Ki3 dominant ACRs (BxK 1,340; OxK 2,280) were few. Over-dominant ACRs were observed in all F1s but were most prevalent in BxO. Taken together, this means that the highly heterotic BxO contains significantly more over-dominant and dominant ACRs than the other two F1 sets (binomial test, p<2.2e^-16^). Moreover, unlike the other two F1s, there were zero BxO under-dominant ACRs. Mirroring the F1 organismal trait over-dominance, the entire BxO genome is as accessible or more than in the parental state. The coincidence of molecular and physiological heterosis suggest that widespread increases in CRE chromatin accessibility contributes to F1 vigour.

The parent-offspring chromatin accessibility comparison classified ACRs based on their genomic position, inter-haplotype homology status and mode of inheritance. We examined the proportional enrichment/depletion of ACRs between these three categories (Figure 2D-F; Data S4), establishing patterns of relatedness between ACR categories that were largely conserved across all three F1s. Haplotype unique ACRs were enriched/depleted for intergenic and TSS locations, respectively. Consistent with the predominance of ‘half-parental’ ACRs, these unique ACRs were also strongly enriched for additive inheritance. In contrast, the high homology, haplotype ‘shared’ ACRs were enriched for TSS locations and depleted for intergenic regions. Lower homology ‘ambiguous’ ACRs were found most often over proximal regions, consistent with intra-specific CRE innovation near, but not at, the TSS. The most pronounced difference between genotypes, was seen at dominant and over-dominant ACRs; BxO dominant and over-dominant ACRs showed the strongest genic ACR enrichments (Figure S3), consistent with dominant and over-dominant accessibility being linked to transcription (Cusanovich *et al*., 2018; Marand *et al*., 2021; Tu *et al*., 2022). The BxO dominant ACRs, were also enriched for low homology ‘ambiguous’ and proximal ACRs, potentially hinting at dominant TF-by-divergent-proximal-ACR interactions.

### TF binding motif bias within ACR categories and genotypes

Further investigating the F1 ACR complements, we searched for DNA sequence motif enrichment/depletion within the ACR categories, but excluded genic ACRs, as they have codon biased sequences, and the exceedingly rare under-dominant ACRs (BxK: eight ACRs; BxO: zero ACRs; OxK: 21 ACRs). Again, despite large intraspecific divergence, the strongest trends were consistent across all genotypes (Figure 3A-D; Figure S4; Figure S5; Data S5). The strongest motif enrichments were seen reciprocally between intergenic and TSS ACRs (Figure 3B-D), with more motifs being prevalent in intergenic ACRs than TSS ACRs. Specifically, many AP2/EREBP motifs were TSS enriched, whereas B3 domain, WRKY, DOF, TCP and other motifs were enriched within intergenic ACRs. Haplotype shared and unique ACRs also had strong and genotype consistent motif enrichments. For example, C4-zinc finger GATA (MA1325.2), MYB (MA1175.2) and G2-like1 (MA1827.2) motifs were reciprocally enriched and depleted in unique and shared ACRs, respectively. Contrastingly, motifs including SPL (MA1056.2), bZIP (MA1822.2) and NAC (MA2036.2) were instead depleted and prevalent in unique and shared ACRs, respectively. Taken together, these genotype conserved enrichments suggest that: I) maize TSS-linked and intergenic CREs are partitioned and II) maize intra-specific CRE innovation contains biased motif variation.

**Figure 3.**
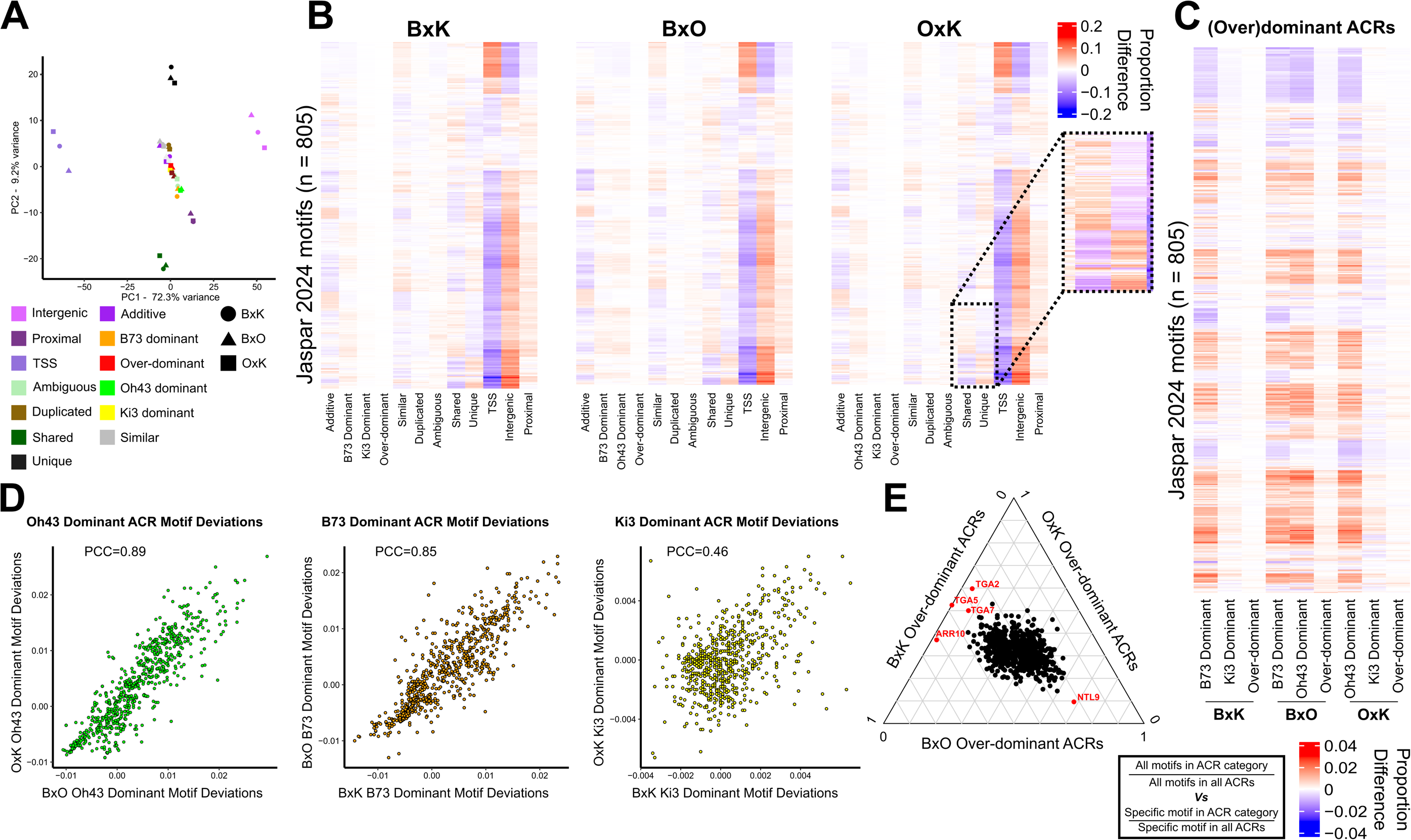
Transcription factor binding motif enrichment/depletion within ACR categories is genotypically stable. (**A**) PCA of all proportional motif enrichments/depletions by ACR category indicate TSS and intergenic ACRs differ the most in motif content, followed by motif differentiation between haplotype shared and unique ACRs. Each of the three dots in a category represent the ACR complement from either BxK, BxO or OxK data, with genotypes (Circle = BxK, Triangle = BxO, Square = OxK) clustering well and exhibiting no consistent trend in PC1 and PC2. Heat maps of individual motif enrichment/depletion in (**B**) BxK, BxO and OxK likewise show strong genotypic similarities. These heatmaps highlight the largely reciprocal TSS and intergenic ACR motif enrichment patterns, and to a lesser extent, motif content reciprocity between shared and unique ACRs. Shared and unique ACRs (inset dashed box – heat colour saturation increased for visualization) do not simply mirror the TSS and intergenic motif relationships, but rather exhibit a distinct enrichment/depletion pattern, just at a reduced magnitude. (**C**) Heat maps of just (over)dominant ACR inheritance, highlight the stability of dominant motif enrichments regardless of hybridization partner and that over-dominant ACRs exhibit weak and F1 variable motif enrichments. (**D**) Haplotype dominant motif enrichment/depletion correlations between F1s illustrate strong consistency in Oh43 and B73, but both weaker (note axes scale) motif bias and F1 correlation over Ki3 dominant ACRs. (**E**) Ternary plot of all three F1 over-dominant ACR motif enrichments/depletions highlight poor, and largely consistent, motif bias in this category. The largest over-dominant motif deviations occurred in BxO, with TGA2/5/7 (SPL-like) and ARR10 motif depletion and NTL9 (NAC) motif enrichments. NTL9 was likewise enriched in BxK. All enrichment/depletion analysis (As outlined in the solid black box) compared the rate of motif occurrence in an ACR category to the rate of motif occurrence in that category in the global ACR set. For example, the proportion of G2-like1 (MA1827.2) motifs in BxO unique ACRs was compared to the proportion of all motifs found in BxO unique ACRs.

Hybrid ACR inheritance categories also had motif enrichment/depletion, but to a lesser magnitude than positional or homology categories and with more genotypic variation (Data S5). Between F1s, dominant ACRs had largely conserved motif enrichments regardless of hybridization partner (Figure 3D); all dominant ACRs exhibited SPL (e.g. MA2452.1) motif enrichments, whereas B73 dominance was enriched for B3-domain (MA0565.3) motifs, Oh43 dominance DOF (MA0977.1) motifs and Ki3 dominance bZIP motifs (MA1822.1). Ki3 haplotype dominance exhibited weaker, and less consistent (Figure 3D), motif enrichments than both other haplotypes, perhaps reflecting the reduced number of Ki3 dominant ACRs. Over-dominant ACRs had the poorest motifs enrichments amongst heritability groups suggesting weaker associations with specific TF families (Figure 3 B and C). Over-dominant ACRs had variable F1 motif enrichments but were weakly enriched for MADS-family motifs (MA0548.3-AGL15, MA1199.1-AGL16, MA0555.2-SVP, MA0005.3-AG) in all F1s (Data S5). BxO over-dominant ACRs had depletion of TGA1 (SPL-like) motifs and both BxK and BxO over-dominant ACRs exhibited NTL9 (NAC) motif enrichment (Figure 3E). BxO over-dominant ACRs also harboured a greater number of stronger motif enrichments than either other F1s (Figure S5; t test; BxO vs BxK, p=1.612e^-12^; BxO vs OxK p=6.97e^-10^), perhaps hinting at more complicated over-dominant regulation in BxO. In summation, BxO heterosis contained more dominant and over-dominant chromatin accessibility than either other F1– and BxO dominance had stronger CRE associations than the over-dominance.

### Narrow-sense heritability estimates of F1 ACRs

The relative influence of environment and genetics on chromatin accessibility is not well understood. These data provide an opportunity to generate h^2^ estimates for chromatin accessibility at each ACR individually by measuring how well the mid-parental value correlates to the observed F1 chromatin accessibilities under controlled growth conditions. Although heuristic, h^2^ calculation here is supported by the preponderance of additive ACR inheritance. To directly compare the independent ATAC-seq datasets, F1 h^2^ was calculated for the B73-reference ACRs from Marand *et al.,* (2025) (Figure 4A). We observed strong overlap between ACRs with high h^2^ and mapped chromatin accessibility quantitative trait loci (caQTL; Fig 4A), indicating that the ACRs under strong genetic control in the three genotypes represent those likewise controlled in the larger population.

**Figure 4.**
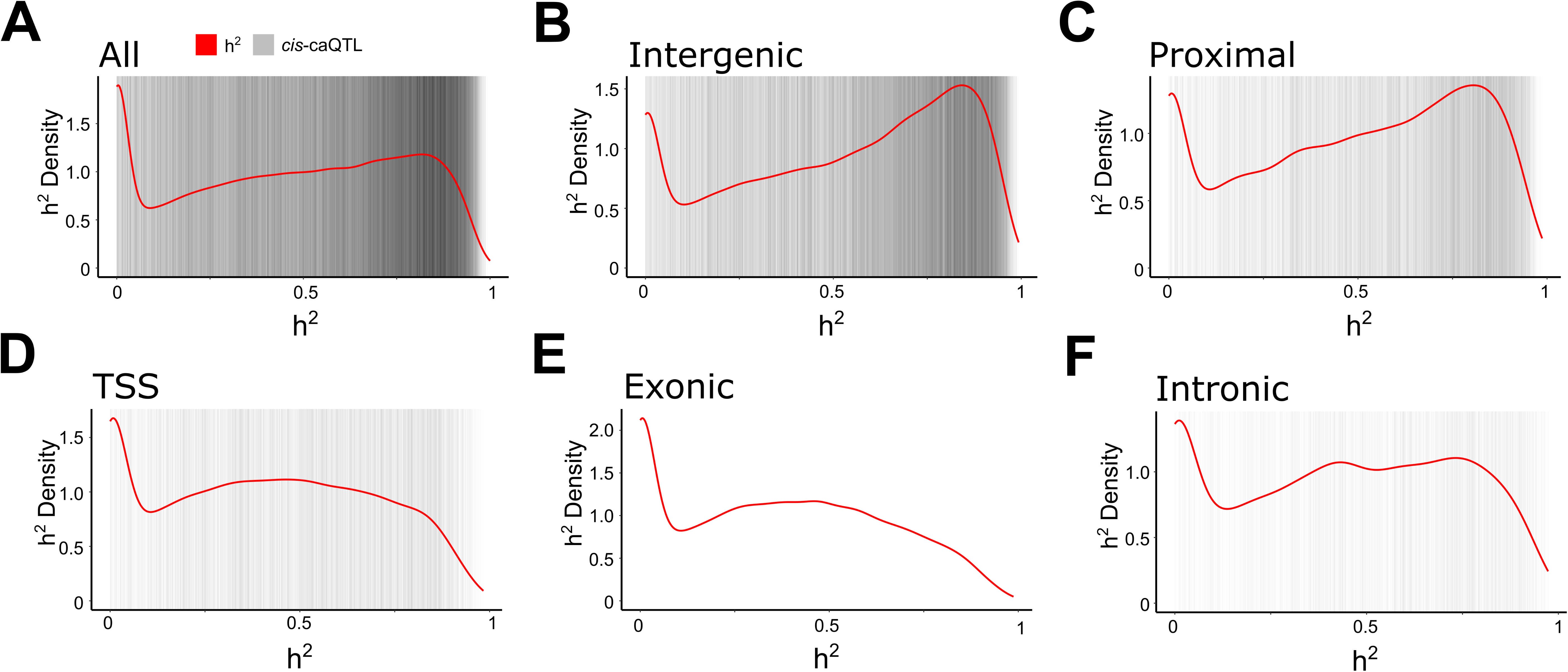
Narrow-sense (h^2^) heritability of chromatin accessibility. (**A**) h^2^ (red line) across all ACRs was relatively high, indicating that additive genetic control of chromatin accessibility is prevalent. These high h^2^ ACRs coincided well with previously detected *cis*-caQTL (grey vertical bars). However, some ACR exhibit almost entire environmental control (h^2^ ∼0), despite being grown under controlled conditions. Split by genomic position, (**B**) intergenic and (**C**) gene proximal ACRs had the highest heritability, with an h^2^ peak of ∼0.85. In contrast, regions near transcription like TSS (**D**) or exonic (**E**) ACRs had lower narrow-sense heritability, with a h^2^ peak closer to 0.5. (**F**) Intronic ACRs had heritability peaks that matched both transcribed ACRs and strictly regulatory ACRs. **Note**: the exonic ACR plot has no *cis*-caQTL hits as they were deliberately excluded from the caQTL analysis.

Further interrogating ACR h^2^, we subcategorized the ACRs based upon their location relative to genes (Figure 4B-F). Across all ACR classes, there were some very low h^2^ ACRs. These low h^2^ ACRs are linked to genes with gene ontology (GO) terms from dynamic cellular processes (Table S3), exemplified by ER-associated misfolded protein catabolic process (GO:0071712), pyridine nucleotide salvage (GO:0019365) and NAD biosynthetic process (GO:0009435). Investigating the function of genes linked to the more heritable ACRs revealed aspects of metabolism like hexose phosphate transport (GO:0015712) and xylan catabolic process (GO:0045493), as well as developmental genes like those involved in regulation of leaf morphogenesis (GO:1901371) and brassinosteroid mediated signaling (GO:0009742). Furthermore, examining higher h^2^ ACRs revealed dichotomous ACR inheritance (Figure 4B-F), where transcribed ACRs (TSS, exonic, intronic) had lower h^2^ than strictly regulatory ACRs (proximal, intergenic). Transcribed h^2^ estimates mirrors those for maize gene transcription (Sun *et al*., 2023), suggesting that chromatin accessibility over transcribed ACRs is largely controlled by RNA polymerase II nucleosome displacement. Moreover, as strictly regulatory ACRs had higher heritability than transcribed ACRs (Figure S6), they are less environmentally sensitive than transcription or transcribed ACRs. Intronic ACR h^2^ appeared different than the other transcribed ACR categories, with two high h^2^ heritability peaks, mirroring both the transcribed and strictly regulatory ACR h^2^ peaks. Given that all introns are transcribed, the prevalence of regulatory h^2^ values was surprising and hints at unique intronic CRE chromatin dynamics. These investigations reveal that proximity to transcription influences chromatin accessibility h^2^. Additionally, under fixed growing conditions, low h^2^ ACRs associated with highly dynamic processes, like cellular homeostasis, while higher h^2^ ACRs likely control more ‘hardwired’ processes like organogenesis and nutrient/carbon utilization.

### Detecting F1 differentially *trans*-regulated ACRs

Returning to the concatenated-reference-based ACRs, we used the parent-offspring measurements to discover F1 ACRs under differential *trans*-regulation. Since F1s exactly preserve the *cis*-regulatory environment, if ACR *trans*-regulation does not differ between parents, F1 chromatin accessibility should simply reflect the change in allele dose, exhibiting ‘half-parental’ chromatin accessibility. However, if inter-parental *trans*-regulatory differences affect an allele, F1 chromatin accessibility will differ from this half-parental value (Figure 5A). To control for haplotype CRE variation, this approach needs allele-specific chromatin accessibility measurements.

**Figure 5.**
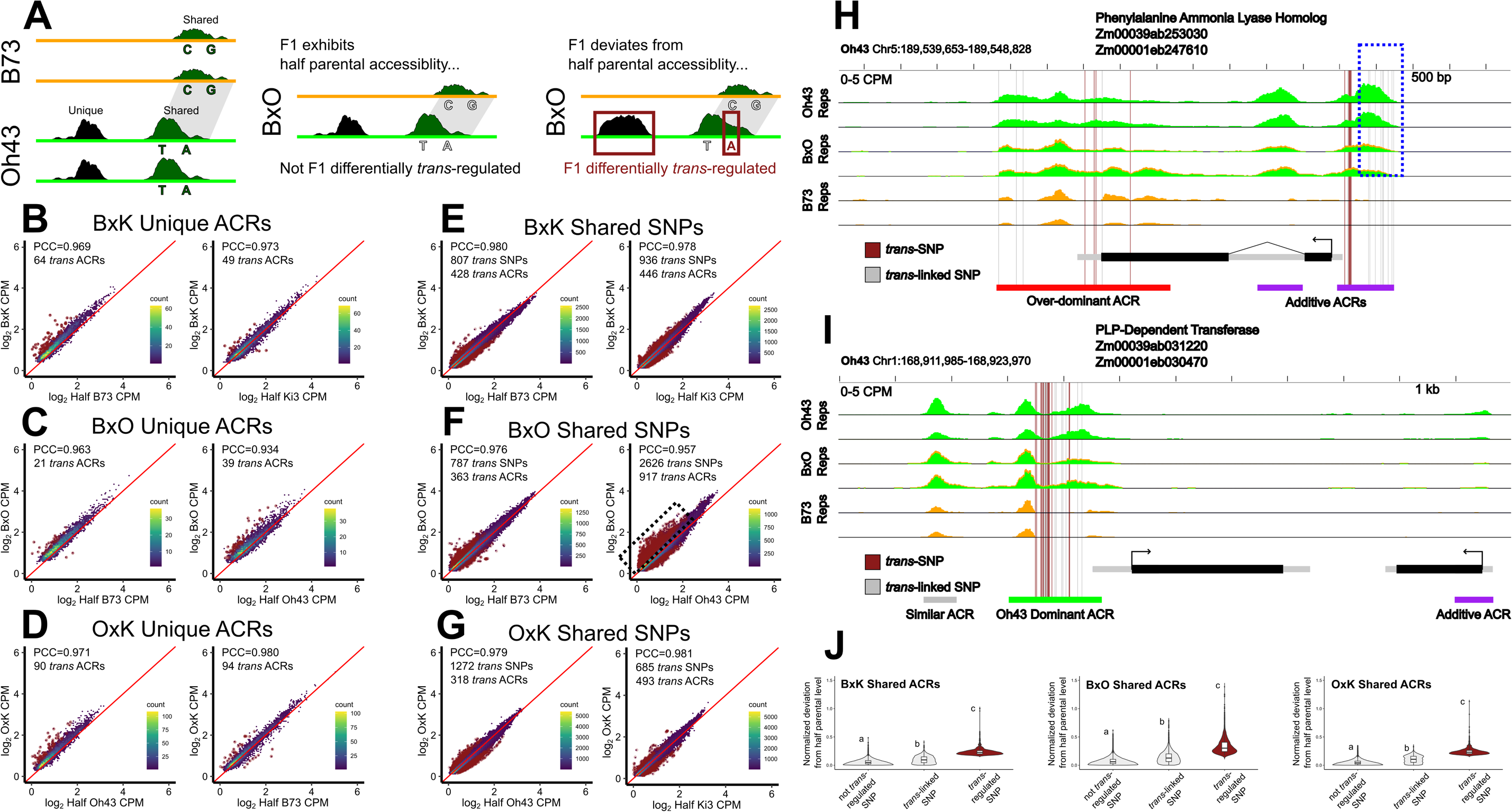
Detection F1 differentially *trans*-regulated ACRs by comparison to allele-dose drive chromatin accessibility levels. (**A**) Exploiting the chromosome-resolved nature of both haplotype unique ACRs (black ACRs) and SNP-containing tn5 fragments from shared ACRs (dark green ACRs), we looked for signs of F1 differential *trans*-regulation. If F1 accessibility is unaffected by the novel *trans*-regulatory environment, then chromosome-resolved F1 accessibility should reflect the change in allele dose and resemble half the accessibility seen in the inbred parent (light grey ACRs/SNPs; not F1 differentially *trans*-regulated). However, if chromosome-resolved F1 accessibility deviates from that predicted by allele dose, it indicates novel F1 *trans*-regulation (burgundy ACRs/SNPs; F1 differentially *trans*-regulated). Examining unique ACRs in (**B**) BxK, (**C**) BxO and (**D**) OxK reveal that most resemble the half parental level, supporting strong *cis*-control of unique ACR accessibility. However, regardless of the heterotic strength, F1 differentially *trans*-regulated ACRs were apparent in all groups (burgundy points). Red line depicts half parental accessibility. Examining SNPs in shared ACRs in (**E**) BxK, (**F**) BxO and (**G**) OxK likewise highlighted the strong *cis*-control of chromosome-resolved accessibility, while revealing F1 differentially *trans*-regulated SNPs. Here, however, the most heterotic BxO F1 exhibited more F1 differential *trans*-regulation, which was especially pronounced over the Oh43 haplotype (dashed black box). Red line depicts half parental accessibility. (**H,I**) Two genome browser shots of the Oh43 haplotype (chromosome) depicting example genes with linked F1 differentially *trans*-regulated SNP (burgundy vertical lines), and *trans*-linked SNPs (grey vertical lines). Two biological replicates are shown for each genotype, with the B73 inbred samples serving to illustrate regions where inter-haplotype mapping occurs, highlighting the need to use SNPs, not coverage, to obtain allele-specific chromatin accessibility measurements. Tn5 fragments containing the *trans*-regulated SNPs differed (greater than two standard deviations higher than the population mean) from the predicted ‘half-parental’ value that is consistent with the F1 change in inbred allele dose. Most ACRs and SNPs exhibited ‘half-parental’ accessibility, as exemplified in the blue dashed box (in panel H). Note the absence of cross mapping from the B73 haplotype in the blue dashed region, justifying the approximation of coverage to allele-specific accessibility. Our stringent *trans*-regulation thresholds likely cause false negatives, as exemplified by the tight linkage between some F1 differentially *trans*-regulated SNPs and *trans*-linked SNPs – as such, all subsequent work was done on shared ACRs with any F1 differentially *trans*-regulated SNP. (**J**) Shared ACRs with F1 differentially *trans*-regulated SNPs, often harbored other SNPs, dubbed *trans*-linked SNPs. These *trans*-linked SNPs had a much-attenuated deviation from the expected half parental value across all genotypes, stressing that F1 differential *trans*-regulation may only involve a subset of an ACR’s CRE complement. Letters denote statistically similar groups, as determined by one-way ANOVA with post hoc Tukey’s HSD test (all comparisons p<2.2e^-16^).

Haplotype unique ACRs represent a good source of allele-specific chromatin accessibility. To ensure these unique ACRs were not an artefact of peak calling, we excluded unique ACRs with ATAC-seq reads originating from the other inbred haplotype. Comparison of F1 unique ACR chromatin accessibility to half the parental inbred value (Figure 5B-D) revealed very high correlations (Pearson Correlation Coefficient; PCC 0.898-0.978), indicating that chromatin accessibility at haplotype unique ACRs is largely driven by *cis* effects. However, some unique ACRs deviated from the *cis*-regulation prediction. To stringently identify unique ACRs with F1 differential *trans*-regulation, we selected ACRs that, on average and across n-1 replicates, deviated the expected ‘half-parental’ values (Figure S7), revealing 357 unique ACRs with F1 differential *trans*-regulation (Data S1,2,3).

Another measure of chromosome-specific chromatin accessibility is Tn5 fragments containing single nucleotide polymorphisms (SNPs) that delineate their haplotype origin. Extracting allele-specific chromatin accessibility is complicated by the concatenated-reference approach, as a single read can map to both haplotypes. Therefore, to extract accurate allele-specific chromatin accessibility, we aligned the shared ACR sequences from parental haplotypes, matched reference/alternative SNP pairs and then quantified reads aligning to this region with the appropriate SNP (Figure 5 EFG). As expected, most SNPs in shared ACRs had equivalent allele-specific reads, reflecting similar chromatin accessibility values in both parents (Figure S8). As with the unique ACRs, the F1 shared ACR allele-specific reads were strongly correlated to the halved parental value (PCC; 0.957-0.981), indicating strong *cis*-control. However, some allele-specific reads deviated from their expected values, with 7,113 SNPs, across 2,965 ACRs, exhibiting F1 differential *trans*-regulation (Figure 5E-G). Examining genome browser shots of genes with linked F1 differential *trans*-regulated SNP (two examples shown Figure 5H and I) highlighted two observations: i) some ACRs show clustered F1 differential *trans*-regulated SNPs, suggesting F1 modular CRE chromatin accessibility and/or modified RNA polymerase II nucleosome displacement and ii) other ACRs have SNPs tightly linked (*trans*-linked SNPs) to F1 differential *trans*-regulated SNPs, that fail to pass our conservative threshold, suggesting our strict approach results in false negatives at the SNP level. Specifically, these 2,965 ACRs harbour 35,597 *trans*-linked SNPs that did not pass the stringent filter for F1 differential *trans*-regulation. These *trans*-linked SNPs deviated from the expected half-parental value less than *trans*-regulated SNPs, but more than SNPs in ACRs without F1 differential *trans*-regulation (Fig. 5J). As such, we identified any shared ACR with at least one F1 differential *trans*-regulated SNP as an F1 differential *trans*-regulated ACR.

Our study design places each haplotype with two different F1 *trans*-regulatory partners, and one combination, BxO (stiff stalk x non-stiff stalk), causes strong agronomic heterosis. To better understand the genetic underpinnings of this strong heterosis, B73 and Oh43 haplotype-resolved ACR *trans*-regulation was compared between F1s. B73 and Oh43 had unique ACRs differentially *trans*-regulated at lower rates with the strong heterotic partner (B73;BxO lower than BxK; binomial test; p=5.2e^-7^; Oh43;BxO lower than OxK; binomial test; p=1.1e^-3^), suggesting unique ACR *trans*-regulation is independent of heterotic strength. However, shared ACR *trans*-regulation appeared more related to good agronomic heterosis. Specifically, BxO shared ACRs on the Oh43 haplotype displayed a pronounced association with heterosis; *trans*-regulatory differences over Oh43 shared ACRs were dramatically more abundant (BxO higher than OxK; binomial test; p<2.2e^-16^), and more biased towards increased accessibility than when paired with OxK (Figure 5FG). In contrast, the B73 haplotype exhibited similar numbers of *trans*-regulated shared ACRs in BxO and BxK (binomial test; p=0.832). Although less pronounced than the Oh43 haplotype, the B73 haplotype appeared more often *trans*-regulated towards higher chromatin accessibility in BxO than in BxK (Figure 5EF). Our triad of parent-offspring measures indicate the importance of increased chromatin accessibility over haplotype-conserved ACRs, and that the B73 (stiff stalk) genome disproportionately donates *trans*-regulators, lopsidedly upregulating the chromatin accessibility of shared Oh43 (non-stiff stalk) ACRs in BxO seedlings.

Next, we investigated ACR genomic position, inter-haplotype homology status and mode of inheritance representation within these 2,965 ACRs with F1 differential *trans*-regulation (Figure S9). Expectedly, across all F1s, differential *trans*-regulated ACRs were depleted for similar parent-offspring chromatin accessibility levels and were enriched for dominant and over-dominant inheritance (Data S4). Irrespective of genotype, proportionally more haplotype shared ACRs exhibited F1 differential *trans*-regulation than haplotype unique ACRs (binomial test; BxK, p<2.2e^-16^; BxO, p<2.2e^-16^; OxK, p=6.5e^-3^). However, genotypic differences were observed for the genomic position of F1 differential *trans*-regulation; BxO alone exhibited significant enrichment for both genic and intergenic ACRs (binomial test; genic: FDR adjusted p=3.69e^-6^; intergenic: FDR adjusted p=3.4e^-3^). We also examined motif enrichment/depletion within these F1 differentially *trans*-regulated ACRs. Here, TF motif enrichments were modest (Figure S10) and genotypic differences pronounced (DataS5) – SPL motifs were most consistently enriched, but the specific SPL motif and enrichment varied genotypically. In the heterotic BxO F1, the AT-hook (MA2374.1) motif had the strongest overrepresentation. Other motifs also enriched in BxO differentially *trans*-regulated ACRs, included HD-ZIP homeodomain motifs (MA1375.2, MA2344.1, MA1369.2), CCA1(MA0972.1), SPL8 (MA0578.1) and a NAC (MA2045.2) motif. These enrichments suggest that ACRs under F1 differential *trans*-regulation largely coincide with dominant and over-dominant ACRs and that *trans*-regulated TF motif bias, although weak, varied genotypically.

### Combining parent-offspring measurements with a diversity panel to detect *trans*-caQTL

Parent-offspring chromatin accessibility measurements establish which ACRs are differentially *trans*-regulated in the offspring but cannot reveal relationships between variable *trans*-regulators and ACRs. However, the association between genetic variation and unlinked effects can be established using diversity panels. Here, we used an environmentally and developmentally matched caQTL diversity panel(Marand *et al*., 2025) to map *trans*-regulatory variants. However, *trans*-caQTL detection is often occluded by small effect sizes and the immense multiple testing burden arising from associating all ACR by all genetic variants – our caQTL panel tested 82,098 non-genic ACRs by 481,944 intra-ACR SNPs, which, after removing linked variants (<5Mbp), left 39,295,217,450 associations. With so many associations, only 322 *trans* SNP-by-ACR interactions pass Benjamini & Hochberg (Benjamini and Hochberg, 1995) false discovery rate correction (FDR = 0.05; Table S4). However, independent filtering(Bourgon *et al*., 2010) can reduce the testing burden by using orthogonal data to subset the variants or ACRs tested. Given the inclusion of B73, Oh43 and Ki3 in the diversity panel, the environmental and developmental matching, and the concordance between the h^2^ estimates and *cis*-caQTL, we reasoned that F1 differentially *trans*-regulated ACRs, would be enriched for *trans* SNP associations in the larger population; that is, only examining ACRs exhibiting F1 differential *trans*-regulation might bolster *trans*-variant associations in the diversity panel.

To best integrate with the B73-reference based caQTL, we only considered BxK or BxO differentially *trans*-regulated ACRs, which directly correspond to a B73 ACR; of the 108,843 ACRs detected in Marand *et al*., (2025), 2,008 and 3,758 ACRs intersected the BxK and BxO differentially *trans*-regulated ACRs, respectively. This independent filter reduced the number of tests considered to 728,926,212 and 1,311,136,899 for BxK and BxO, respectively. For the respective BxK and BxO filters, 31 and 26 *trans* SNP-by-ACR passed FDR=0.05 correction (Table S5), representing a ∼5.2 (BxO) and ∼2.4 (BxK) fold higher discovery rate compared to the unfiltered ACR set (test of one proportion; BxK: p<2.2e^-16^; BxO: p=3.25e^-6^). Additionally, all *trans* SNP-by-ACR relationships found after independent filtering were new interactions missed in the 322 SNP-by-ACR relationships discovered with the unfiltered ACR set. The 31 and 26 *trans* SNP-by-ACR relationships represent all those found in the 172 genotypes measured in the Marand *et al*., (2025); filtering for SNPs variable in B73, Oh43 and Ki3, left only 1 *trans*-associated variant in each F1 parental combination (BxK: chr2:189030540; BxO: chr10:149076828). The BxK *trans*-variant was linked to a Sterile alpha motif (SAM) domain gene (Zm00001eb100110) and *ZIM-transcription factor13* (Zm00001eb100130). The BxO variant was between an RNA binding protein (Zm00001eb432880) and an uncharacterized gene (Zm00001eb432890). Although independent filtering based on BxK or BxO differentially *trans*-regulated ACRs provides more sensitivity compared to the entire ACR set, this approach remained limited in discovery potential, unveiling few *trans*-regulatory relationships, and highlighting the need for better methodologies.

### Multiple discovery filtering (MDF): a new *trans*-regulatory variant detection approach

There is a fundamental disconnect between the number of *trans*-regulators mapped in structured populations and multi-parent diversity panels. Bi-parental studies uncover numerous, small-effect-size, *trans*-relationships (Brem *et al*., 2002; Scheetz *et al*., 2006; West *et al*., 2007; Hasin-Brumshtein *et al*., 2016; Parker *et al*., 2024) but multi-parental panels identify few with confidence due to larger testing burdens and lower minor allele frequencies (MAF) depressing power. Bi-parental studies indicate that *trans*-acting variants regulate many unlinked loci (Brem *et al*., 2002; Scheetz *et al*., 2006; West *et al*., 2007; Hasin-Brumshtein *et al*., 2016; Parker *et al*., 2024), and this ‘hotspot’ trend extends to multi-parent populations (Morloy *et al*., 2004; Dixon *et al*., 2007; Võsa *et al*., 2021). Knowing this relationship, we devised a novel multiple test correction solution that focuses on confidently identifying *trans*-regulatory variants that associate with many loci. This pattern mirrors the *trans*-regulatory patterns established in structured populations, revealing variants linked to altered *trans*-regulators with widespread genomic effects.

To evaluate the effectiveness of this MDF approach, we set out to re-discover *trans*-expression-QTL (eQTL) hotspots found by others. Using a well-powered *Mus musculus* (mouse) hypothalamus eQTL dataset (Hasin-Brumshtein *et al*., 2016), we masked (artificially increased) their Bonferroni-correction passing p-values before applying MDF and attempted to recapitulate their discovered eQTL hotspots (Figure 6A). Despite significant residual population structure, MDF found both known *trans*-eQTL hotspots (*Mmu*chr1: 171039894-176205262; *Mmu*chr15:96181065-98130818) while uncovering two more *trans*-eQTL regions: *Mmu*chr2:102764753-103924035 and *Mmu*chr4:81759048-89433421. Other potential hotspots may exist but could not be confidently distinguished from population structure noise. Furthermore, filtering (99.9^th^ percentile) variants for the most *trans*-relationships revealed one SNP (rs30718063) at *Mmu*chr1:114457498 with 2544 gene relationships – more than double that found in the previously identified hotspots (1141 genes chr15; 842 genes chr1). Rs30718063 is rare in the sampled Hybrid Mouse Diversity Panel Group (HMDP) population (Bogue *et al*., 2020) (Table S6), precluding detection in the initial study. Supporting rs30718063 biological importance, this SNP is a hypothalamus *cis*-eQTL(Hasin-Brumshtein *et al*., 2016) for *CONTACTIN-ASSOCIATEDPROTEINLIKE5a* (*CNTNAP5a*; ENSMUST00000043725.8), a NEUREXIN-family protein expressed in the developing brain and the hypothalamus(Zeisel *et al*., 2018) with predicted roles in brain development (Gomez *et al*., 2021). Supporting MDF validity and power, we re-capitulated known *trans*-eQTL hotspots, revealed new weaker *trans*-eQTL hotspots, and uncovered a rare and impactful SNP despite analysing masked data.

**Figure 6.**
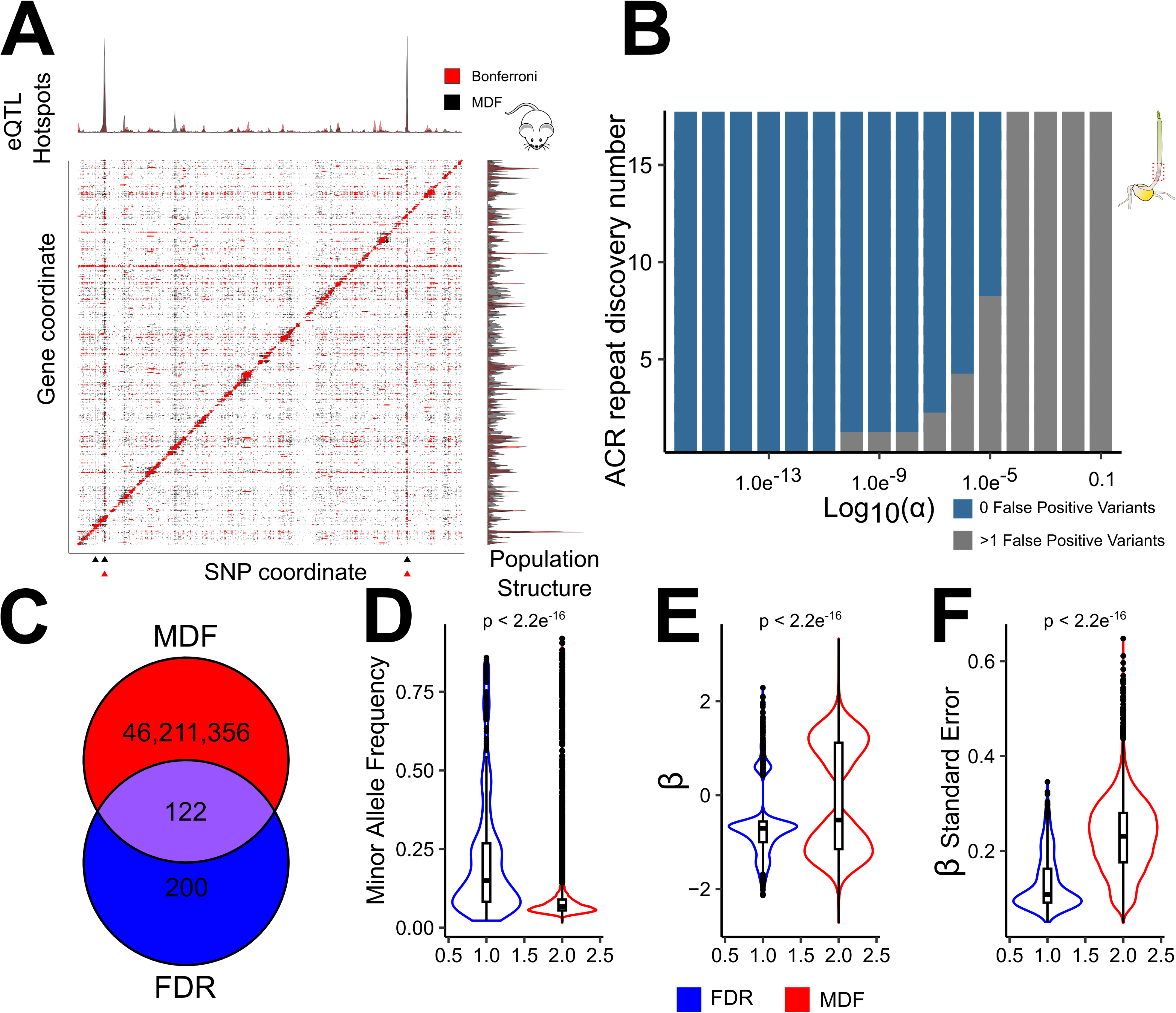
Multiple discovery filtering (MDF) recapitulates known mouse *trans*-eQTL and empowers maize *trans*-caQTL detection. (**A**) To test the validity of our MDF approach, we attempted to repeat the *trans*-eQTL hotspot detection from masked *Mus musculus* (mouse) hypothalamus eQTL data. Despite residual population structure (y-axis histograms), MDF re-discovered the known *trans*-eQTL hotspots (stacked black and red triangles) while supporting at least two novel trans-eQTL hotspots (black peaks x-axis histograms). Moreover, MDF uncovered a rare variant (single black triangle) that both *trans*-associated with many gene expression changes and was a *cis*-eQTL of a *NEUREXIN-family* gene, a known neural development regulator, in the initial study. (**B**) For MDF empowered *trans*-caQTL mapping, we iteratively tuned ACR repeat discovery number (n-DISC) and α thresholds to achieve a false discovery rate comparable to 0.05, which is used traditionally via Benjamini & Hochberg FDR adjustment. (**C**) MDF provides support for many more SNP-by-ACR association than that supported by Benjamini & Hochberg FDR adjustment, although significant overlap between the two SNP-by-ACR association sets exists. Comparing the Benjamini & Hochberg FDR supported and MDF supported *trans*-regulatory variants highlight that MDF (**D**) unveils rarer alleles, (**E**) finds alleles with both larger effect sizes (absolute β) and less bias towards negative β and (**F**) finds alleles with more variable effects in the 172-genotype population. p-values reflect those from a Wilcoxon rank sum (Mann-Whitney U) tests.

To further compare the discoveries of MDF filtering to traditional FDR testing, we returned to the 172-genotype maize caQTL panel (Marand *et al*., 2025). Reassuringly, both FDR correction and MDF with a n-DISC of one produce a near equivalent p-value cutoff (FDR: 1.68e^-12^>p>1.53e^-12^; MDF: 1.0e^-11^>p>1.0e^-12^) at which we expect SNP false positives to round to zero. Iteratively tuning n-DISC and α thresholds (Figure 6B), we achieved a false positive rate near 0.05 (FDR 0.024; Table S7) and revealed 31,612 *trans*-regulatory variants supported by 46,211,478 *trans* SNP-by-ACR interactions. 122 of the 322 SNP-by-ACR interactions that passed traditional FDR corrections (Figure 6C), were rediscovered in this MDF set, significantly higher than the overlap expected due to chance (test of one proportion; p<2.2e^-16^). Compared to FDR testing (Figure 6D-F), MDF found *trans*-regulatory variants at lower MAF (Wilcoxon Rank Sum test, p<2.2e^-16^), larger absolute effect sizes (β, Wilcoxon Rank Sum test, p<2.2e^-16^) and larger effect variability (β standard error, Wilcoxon Rank Sum test, p<2.2e^-16^). Moreover, the FDR discoveries were heavily biased towards negative β values (i.e. *trans* SNP-by-ACR interactions reducing chromatin accessibility), whereas MDF discoveries were better distributed between negative and positive ACR relationships (Figure 6E). MDF *trans*-regulatory variants overlapped with Genome Wide Association (GWAS) SNPs (Wallace *et al*., 2014; Liu *et al*., 2023) at a lower rate (0.76 times lower; test of one proportion; p=7.3e^-05^) than the remainder of the ACR SNPs (Table S8); this is consistent with the notion that extant *trans*-regulatory variants reflect small effect alleles, and that more population trait variation is detectably over strictly *cis*-regulatory variants.

The genes linked to these MDF passed *trans*-regulatory variants were enriched for DNA transcription (GO:0006355; Fisher’s exact test, p=1.1e^-16^; Table S9) and other executive processes, including hormone response (GO:0009755; Fisher’s exact test, p=1.12e^-12^) and signal transduction (GO:0007165; Fisher’s exact test, p=2.8e^-11^). Moreover, these variants were linked to many classical maize genes including those involved in cell fate (*gl15*, *gl2*, *kan1*, *kn1*, *ns1*, *ocl1*, *pebp1*, *ra1*, *ra2*, *ra3*, *si1*, *tan1*, *tb1*, *ts1*, *zag1*), light and hormone signalling (*D8*, *D9*, *phyA1*, *phyB2*, *phyC1*, *phyC2*) and metabolism (*amya3*, *cesa1*, *cesa3*, *cesa6*, *ivr1*, *ivr2*, *sbe1*, *ss1*, *ssu1*, *su2*, *sus1*). Two *trans*-regulatory SNPs (chr1:272262485, chr1:272266500) were found within the *tb1* control region that coordinates *tb1* expression(Studer *et al*., 2011). The ACRs associated with these two *trans*-regulatory variants exhibited mild enrichment for TB1 ChIP-seq (Dong et al., 2019) peaks (one-sided binomial test; chr1:272262485 p=0.052; chr1:272266500 p=0.028) and *de novo* motif discovery(Bailey, 2021) found GGNCCC(N) motif enrichment in both *trans*-associated ACR sets which resembles the core TB1 binding motif (Kosugi and Ohashi, 2002; Dong et al., 2019). Likewise, motif enrichment(Bailey and Grant, 2021) supported TB1 motif enrichment in the ACRs *trans*-associated with both SNPs (MA1430.2; p=2.37e^-7^). Together, MDF passing *trans*-caQTL variants contain hallmarks of expected chromatin accessibility *trans*-regulators, supporting MDF’s utility in unravelling unlinked genetic-to-chromatin state associations.

The 282 maize diversity panel also has previously published *cis*-eQTL data(Kremling *et al*., 2018). Like *cis*-caQTL analysis, *trans*-eQTL detection has been obfuscated by poor power and the multiple testing burden. To look for *trans*-regulatory variants affecting maize gene expression, we realigned diversity panel mRNA sequencing reads and built associations between 19,584 expressed genes and the same set of 481,944 SNPs before applying MDF. As we tested far fewer genes than ACRs the MDF approach was able to more stringently identify *trans*-regulatory iterations; 172,697 *trans*-regulatory variants were supported by 52,982,436 SNP-to-gene interactions (Table S10), with only ∼6 of these variants expected as false discoveries. To investigate these *trans*-regulatory relationships, we first turned to the characterized maize pigment pathway, specifically to *Yellow endosperm1* (*y1*; Zm00001eb271860), which underpins carotenoid level variation in domesticated maize (Chen *et al*., 2024). Examining 23 *y1*-linked *trans*-regulatory variants (Table S11) revealed *trans*-associations with known pigment genes including *deoxy xylulose synthase3* (*dxs3*; Zm00001eb377300), *phytoene synthase2* (*psy2*; Zm00001eb367770) and *White cap1* (*Wc1*; Zm00001eb402840) and other genes putatively involved in terpene metabolism or the methyl-d-erythritol-4-phosphate (MEP) pathway (Zm00001eb003330, Zm00001eb428470, Zm00001eb288110, Zm00001eb323510, Zm00001eb302370, Zm00001eb224640, Zm00001eb201480, Zm00001eb268530, Zm00001eb270010, Zm00001eb414190, Zm00001eb325560). We also investigated six *trans*-regulatory variants linked to an iron metabolism gene (Zm00001eb377130; Table S12) and lead GWAS-hit for maize chlorophyll content(Ali *et al*., 2025), which revealed *trans*-associations with known chlorophyll biogenesis (*camouflage1*; Zm00001eb233020, Zm00001eb047780, Zm00001eb044650, Zm00001eb431800, Zm00001eb044650, Zm00001eb287860, Zm00001eb199950) and iron electron transport (*ferredoxin3*; Zm00001eb063710, Zm00001eb167930, Zm00001eb295180, Zm00001eb321670, Zm00001eb295180) genes. Together, we see that MDF passing variants are supported by biologically relevant SNP-to-gene relationships, highlighting MDF as a potent tool to identify putative genetic interactions for future molecular characterization.

### Pairing parent-offspring measurements with MDF unveils heterotic *trans*-regulatory variants

To unravel heterotic *trans*-interactions, since MDF detection power scales inversely with the number of tests, we again filtered *trans*-caQTL tests to only include BxK or BxO differentially *trans*-regulated ACRs. Associating all SNPs and these ACR subsets, 36 and 2,102 *trans*-regulatory variants pass MDF for BxK and BxO differentially *trans*-regulated ACRs respectively (Table S12). To investigate if the limited BxK *trans*-regulatory variant discovery was due to the number of ACRs associated, we downsampled the examined BxO ACRs to match that in BxK (2,008 ACRs) and used the BxK n-DISC and α MDF thresholds. This BxO downsampled ACRs set (Table S13) revealed 9,670 *trans*-regulatory variants, indicating that strictly technical factors do not underlie the paucity of BxK *trans*-regulatory variant discoveries; rather, the BxO differentially *trans*-regulated ACRs appear to be controlled by stronger *trans*-regulatory variants in the sampled population. These *trans*-regulatory variants reflect all possible *trans*-regulatory variants in the 172 genotype caQTL panel; filtering these *trans*-regulatory variants to only include genetic differences in BxK and BxO, left 1 and 137 *trans*-regulatory variants respectively. The sole SNP polymorphic in BxK (chr8:134587536) was linked between a metal transporter (Zm00001eb354910) and a bromodomain protein (Zm00001eb354900). However, this single *trans*-regulatory variant reveals little about the biological processes that vary in the BxK F1.

In contrast, the genes linked to the 137 variable BxO SNPs (Table S14) provided insights into the biological processes underpinning vigorous F1 seedling growth. These variants were linked to characterized master regulators, including kinases, metabolic integration points, and TFs (Table 1). BxO *trans*-regulatory variants highlighted one AT-hook TF (Zm00001eb434070) and an SPL-binding TF (Zm00001eb322280), that had their cognate DNA binding motifs enriched within the underlying BxO differentially *trans*-regulated ACRs. The known roles of these regulators are consistent with less conservative seed resource use (see discussion), driving more rapid growth through increased cellular division, carbon utilization and investment in photosynthetic pathways. Beyond highlighting known growth processes, our approach highlights the specific genes/variants behind seedling vigour. This provides putative markers for breeding improved seedling heterosis through extant variation and targets through which editing could improve maize seedling growth more generally.

Each *trans*-regulatory variant that associates with the BxO independent filter, is supported by many individual ACR-by-SNP associations (Fig 7A). The metrics from these associations measure the support for each variant, as well the strength of its genomics impacts. Pairwise comparisons of these metrics (Figure S11), revealed a positive relationship between variant effect size (mean β) and the number of ACR associations (Figure 7BCD; p<2.2e^-16^), suggesting that impactful variants have large chromatin accessibility impacts at many loci at once. Moreover, MAF was inversely correlated with both the numbers of ACRs associated and mean β (Figure 7EF respectively, Pearson’s correlation coefficient=-0.29; p<2.2e^-16^; Pearson’s correlation coefficient=-0.57; p<2.2e^-16^). This mirrors what was observed for maize *cis*-eQTL(Kremling *et al*., 2018), indicating that, at least for this ACR-subset, large effect chromatin accessibility *trans*-regulatory alleles are rare in domesticated maize. To contrast the BxO variable *trans*-regulatory variants to the rest of the diversity panel, we compared the constituent ACR-by-SNP association metrics between the 172 variable BxO SNPs with the 1,940 *trans*-regulatory SNPs that were fixed in the BxO parents. This comparison revealed that BxO *trans*-regulatory variants had weaker effects than the rest of the population (Figure 7G-L; mean β, t-test; p<2.2e^-16^; number of ACRs associated, t-test; p=4.64e^-8^) but were present at higher allele frequency (MAF, t-test; p=1.53e^-8^). This suggests that the BxO *trans*-regulatory variation reflects alleles of attenuated effect sizes that are relatively prevalent in domesticated maize diversity.

**Figure 7.**
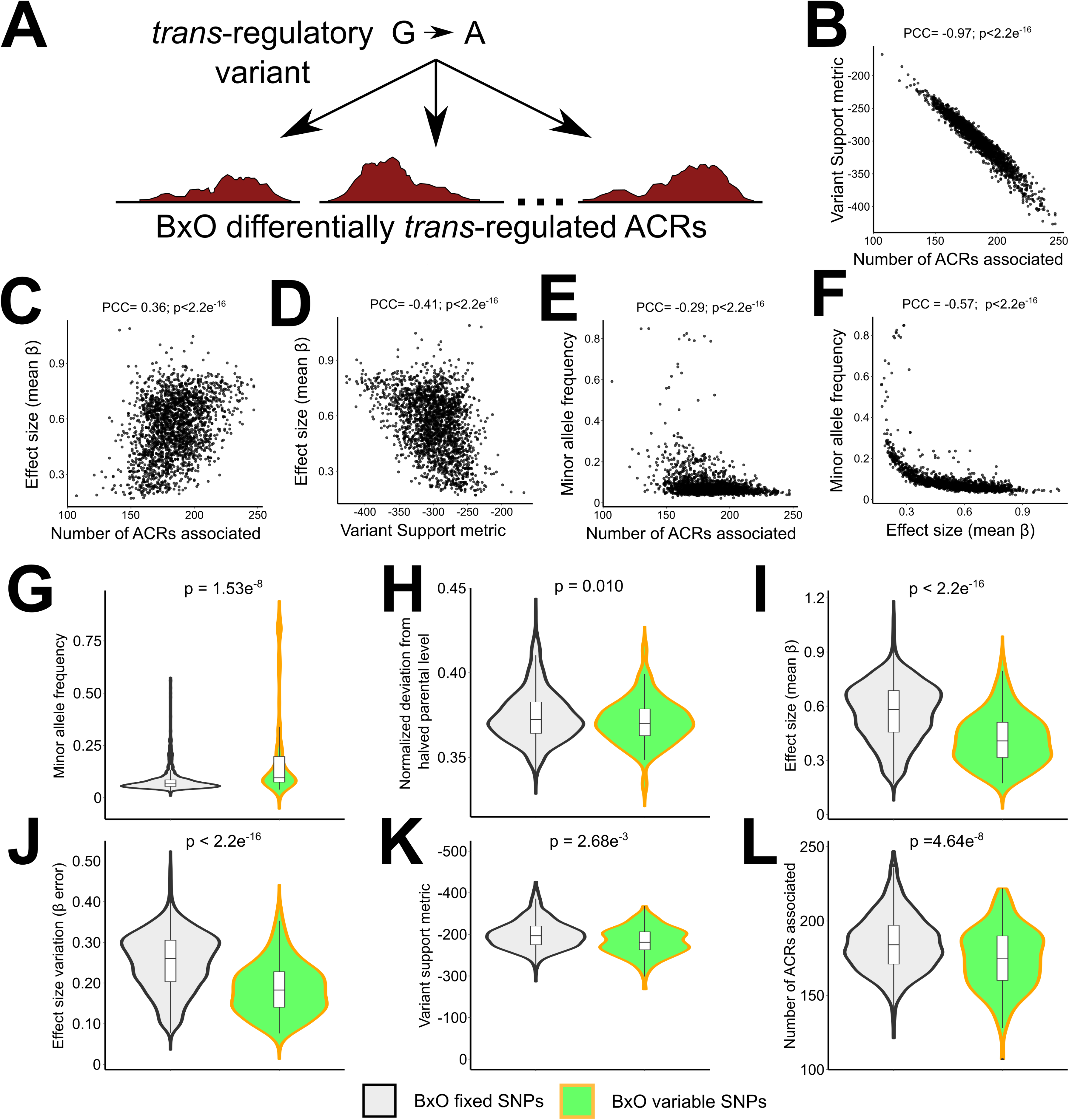
*Trans*-regulatory variants that associate with BxO differentially *trans*-regulated ACRs are more common and have attenuated effects compared to SNPs segregating in other maize accessions. (**A**) MDF intrinsically links many ACRs to a single *trans*-regulatory SNP, providing metrics to compare the effects of *trans*-regulatory variants on the genome. (**B**) Within the *trans*-regulatory variants influencing BxO differentially *trans*-regulated ACRs, the variant support metric (log_10_[probability of a false positive]) correlates strongly with the number of associated ACRs. Effect size (mean absolute β) has a loose relationship with both (**C**) the number of ACR trans-associated and (**D**) the variant support metric. Minor allele frequency (MAF) inversely correlates with both the (**E**) number of *trans*-associated ACRs and the (**F**) effect size (mean absolute β) on those ACRs, indicating that rare alleles have more potent *trans*-regulatory consequences. Comparing the SNPs variable between B73 and Oh43 (BxO variable SNP) and those fixed in both inbred (BxO fixed SNPs) revealed that (**G**) the *trans*-regulatory variation housed in BxO is at a higher MAF than the SNPs in the rest of the population and (**H**,**I**,**J**,**K**,**L**) that BxO *trans*-regulatory variation has smaller effect sizes than that seen in the larger maize population.

## Discussion

Understanding the mechanisms of F1 heterosis has been a perennial challenge. However, it is clear that heterosis is complex, involving interactions between unlinked regions of the genome (Stuber et al., 1992; Kaeppler, 2012; Riedelsheimer et al., 2012; Li et al., 2018; Xiao et al., 2021; Wang et al., 2023a). This F1 *trans*-regulation is thought to cause non-additive F1 transcription and organismal phenotypes. Here, we used bulk seedling ATAC-seq measurements from three inbred-parent-to-F1-offspring combinations, of varying heterotic strength, to study chromatin accessibility inheritance. Using a concatenated reference approach, we comprehensively interrogated haplotype-resolved ACR variation and characterized genomic ACR inheritance. We revealed that CRE chromatin accessibility mirrored (over)dominant F1 plant phenotypes, with heterotic F1 ACRs being as accessible, or more, than either inbred parent. We combined these parent-offspring ATAC-seq measurements with a tissue-matched caQTL mapping population(Marand *et al*., 2025) and a novel MDF approach to interrogate F1 variable *trans*-regulators. Demonstrating MDF utility, we provide insights into the biological processes, and *trans*-regulatory variants, underpinning BxO (stiff stalk x non-stiff stalk) seedling heterosis.

### CRE chromatin accessibility inheritance patterns are genetically controlled with remarkable genotypic consistency

Our paired parent-offspring measurements established the modes of inheritance for individual ACRs. Of ACRs with F1 chromatin accessibility differences, under-dominance was nearly absent, over-dominance was rare, dominance was prevalent, and additive inheritance was most common. Under-dominance paucity is not surprising, as under-dominance associates with hybrid incompatibility, frequently seen in interspecific crosses(Coolon *et al*., 2014; Kitano and Okude, 2024). The abundance of additive inheritance allowed us to make ACR-resolved narrow-sense heritability estimates of chromatin accessibility. Although some ACRs exhibited low genetic control (h^2^ ∼0) under controlled growth, most had a strong relationship with the predicted mid-parent value. Comparison with our tissue and environmentally matched diversity panel(Marand *et al*., 2025) revealed widespread concordance between high h^2^ and *cis*-caQTL detection. Despite only sampling three of the 172 genotypes in the larger population, *cis*-caQTL detection over low h^2^ ACRs, was infrequent. This suggests that the ACRs under genetic control are similar between genotypes, although the underlying mechanism of control likely varies.

Non-transcribed intergenic and proximal ACRs had very high heritability (h^2^ peak ∼0.85) indicating near complete additive genetic control under fixed growth conditions. Many of these non-transcribed ACRs exhibited ‘half parental’ inheritance, exhibiting chromatin accessibility reflecting the halved allele dose. In contrast, transcribed ACR h^2^, which reflects regulatory CRE dynamics and RNA polymerase II nucleosome displacement, was lower than h^2^ over non-transcribed ACRs. Transcription-proximal h^2^ mirrored transcription inheritance, with our transcribed ACR h^2^ peak being nearly identical to maize gene expression H^2^(Sun *et al*., 2023). This suggests that active transcription effectively conceals the nucleosome remodeling processes that maintains chromatin accessibility away from genes. Moreover, the lower h^2^ indicates that transcription-coupled nucleosome eviction is more sensitive to the environment and under less additive genetic control. This latter conclusion is supported by the enrichment between non-additive ACR inheritance, notably (over)dominance, and TSS or genic ACR locations. Framed another way, this indicates that non-additive F1 chromatin accessibility often reflects (over)dominant transcription.

We also examined enrichment/depletion of TF motifs within ACRs of differing genomic position (excluding genic ACRs), inter-haplotype homology status and mode of inheritance (excluding under-dominant ACRs). The strongest motif biases varied by genomic position and were strikingly invariant amongst genotypes. Motif positional bias was predominantly complementary motif enrichment/depletions between intergenic and TSS locations. It has long been appreciated that certain motifs (e.g. TATAA box, initiator motif) are biased relative to the TSS(Haberle and Stark, 2018). Likewise, emerging evidence in compact plant genomes supports diverse TF CRE partitioning by TSS-proximity(Shukla *et al*., 2025). However, in large genomes, repetitive DNA expansion moves CREs from their cognate gene targets(Schmitz *et al*., 2022), producing distal ACRs(Eli *et al*., 2016; Oka *et al*., 2017; Lu *et al*., 2019; Ricci *et al*., 2019). To the best of our knowledge, this is the first report of plant motif partitioning between TSS and intergenic ACRs, although distal enhancer-like regions (DELs) have functional specialization in metazoan genomes(Abascal *et al*., 2020). This, along with the observed h^2^ difference, suggests that transcriptional regulation through distal TF-binding involves molecular mechanisms distinct from those nearest to RNA polymerase II transcription. Since maize distal regulatory regions have demonstrated phenotypic impacts(Stam *et al*., 2002; Salvi *et al*., 2007; Studer *et al*., 2011; Whipple *et al*., 2011; Zheng *et al*., 2015; Liu *et al*., 2015; Eli *et al*., 2016; Huang *et al*., 2017; Lu *et al*., 2019; Ricci *et al*., 2019; Tian *et al*., 2019), future study into the molecular distinctions between intergenic and TSS proximal *cis*-regulation may enhance our mechanistic understanding of trait variation in large repetitive genomes.

We also observed TF motif deviations in ACRs of different inter-haplotype homology status, with haplotype unique and shared ACRs having the strongest biases. These were milder than that observed for genomic position, but, amazingly, were still genotypically consistent. The shared ACRs contain the CREs with the highest intra-specific conservation. Here, genotypic consistency in motif deviation was expected; certain TF regulatory networks may be too integral to vary without maladaptive pleiotropy, restricting cognate CRE innovation over brief evolutionary timescales. In contrast, the consistent TF motif bias in haplotype unique (presence-absence variation) ACRs was very surprising, as we sampled tropical, stiff stalk and non-stiff stalk inbreds, representing divergent phylogenetic branches of domesticated maize(Hu *et al*., 2021). Some of this TF motif bias likely represents the enrichment/depletion of intergenic/TSS ACR (and their motif bias) within unique and shared ACRs, respectively. However, motif bias (Figure 3B-D) does differ between these groups; i.e. intergenic TF motif bias is similar, but not the same, as that within unique ACRs and, likewise, TSS TF motif bias is not identical to that in shared ACRs. Rather, despite large genetic distances, it appears that CRE presence-absence variation follows a consistent pattern in extant maize. The mechanism here is not immediately apparent, but we have two hypotheses: I) historical artificial selection, despite different environments, continually favours CRE gain/loss within a consistent subset of the maize CRE ‘grammar’ and/or II) CRE gain/loss is driven by transposon expansion/contraction, with maize transposon-encoded CREs likewise being restricted in their CRE ‘grammar.’ A more comprehensive study of extant ACR complements, ideally exploiting the available high-quality maize reference genomes(Hufford *et al*., 2021), will hopefully shed more light on this remarkable genotypic consistency.

### Seedling heterosis accompanies non-additive chromatin accessibility inheritance

Heterosis is a polygenic trait, with dominant, over-dominant and epistatic F1 *trans*-interactions summing to produce non-additive organismal phenotypes. Here, we observed parallel chromatin accessibility phenomena, with heterotic F1s exhibiting abundant dominant and over-dominant chromatin accessibility inheritance. This chromatin accessibility (over)dominance was enriched for gene-proximal ACRs, suggesting that F1 (over)dominant transcription is more prevalent than (over)dominant chromatin accessibility over strictly regulatory ACRs. TF motif enrichment was both stronger in dominantly inherited ACRs than in over-dominant regions and largely consistent regardless of hybridization partner. This suggests dominance is TF driven, with the TF complement from a haplotype being sufficient to maintain chromatin accessibility even at a halved allele dose. Over-dominant ACRs had weaker and less consistent motif enrichment, perhaps hinting at more complex, less TF-driven regulation; indeed, the mapped *trans*-regulatory variants often involved kinases and metabolic hubs, which integrate into transcription via many TF-families(Laplante and Sabatini, 2013; Peixoto and Baena-González, 2022). However, over-dominant ACRs were consistently enriched for MADS-family TF motifs, suggesting these TFs, and their gene targets, are relevant to seedling heterosis. Others have implicated *MADS69*, an important floral regulator(Liang *et al*., 2019), in maize heterosis(Xiao *et al*., 2021), but *MADS69* expression is phenologically separate from seedling vigour(Woodhouse *et al*., 2022). Despite the demonstrable agronomic importance of *MADS* genes in domestication(Zhao *et al*., 2011; Liang *et al*., 2019), heterosis(Xiao *et al*., 2021), and bioengineering(Wu *et al*., 2019), we found no *trans*-regulatory variants near *MADS* genes. Rather, we suspect that MADS-TFs may be common targets of the non-TF *trans*-regulators we discovered.

This data set also allowed us to ascertain which shared or unique ACRs exhibited the strongest changes in F1 chromatin accessibility due to *trans*-regulatory differences. For unique ACRs, there seemed to be no relationship between the amount of F1 differential *trans*-regulation and heterosis. In contrast, BxO shared ACRs were more abundantly *trans*-regulated than the less heterotic F1s, suggesting *trans*-regulation of conserved parts of the genome correlates with heterosis. Curiously, we report bias in this F1 differential *trans*-regulation, specifically that the Oh43 (non-stiff stalk) haplotype exhibited dramatic chromatin accessibility increases only when paired with B73 (stiff stalk). This pattern suggests B73 *trans*-regulators elicit increased chromatin accessibility, and presumably transcription, of the Oh43 haplotype. Future work should investigate if this trend is maintained across other stiff stalk x non-stiff stalk F1 seedlings and in other tissues. Motif enrichment of these BxO differentially *trans*-regulated ACRs revealed modest motif enrichment. However, the top enriched motifs largely correspond to Arabidopsis TFs that delineate the meristematic division-differentiation boundary(Talbert *et al*., 1995; Vroemen *et al*., 2003; Horstman *et al*., 2015; Shi *et al*., 2024). Although the cognate maize TFs, and their roles, remain uncertain, the coincidence of Arabidopsis division-differentiation motifs and the mapped cell-division *trans*-regulators suggest that these ACRs may underpin vigorous F1 cell proliferation.

### MDF empowers *trans*-regulatory variant detection providing insights into the underpinnings of seedling heterosis

Genomic *trans*-regulatory relationships possess a striking duality – population variance in *trans*-regulators is very small, yet molecular study repeatedly highlights *trans*-regulators, like TFs, kinases and metabolic hubs, as critical integration points in transcriptional responses(Katagiri and Chua, 1992; Paz-Ares *et al*., 2002; Lambert *et al*., 2018). Indeed, it is postulated that the oversized roles of these genes restrict intra-specific functional variation(Wray *et al*., 2003; Swanson-Wagner *et al*., 2009), by purging novel alleles. Yet the potential to manipulate these large effect *trans*-regulators has been demonstrated in both pharmacological(Attwood *et al*., 2021) and agronomic settings (Nuccio *et al*., 2015; Wu *et al*., 2019; Zhan *et al*., 2025). However, challenges(Swenson *et al*., 2024), and opportunity(Darnell, 2002), remain in developing TF-targeted drugs. Functional manipulation also remains limited by the ability to discover and understand these *trans*-regulatory relationships. GWAS effectively links *cis*-regulatory genetic variation to transcriptional (and other) genomic signals. However, establishing genome-wide *trans*-regulatory relationships remains difficult; constrained functional variation, small effect sizes and an immense multiple testing burden prevents routine *trans*-QTL establishment in diverse populations.

Structured populations, which inflate MAF and statistical power, can establish *trans*-regulatory relationships. Consistent with molecular models, structured populations demonstrate that one *trans*-regulatory variant influences many genes simultaneously, forming *trans*-QTL ‘hotspots’(Brem *et al*., 2002; Scheetz *et al*., 2006; West *et al*., 2007; Hasin-Brumshtein *et al*., 2016; Parker *et al*., 2024). Here, we developed a new approach, MDF, which finds *trans*-regulatory variants in diversity panels by looking for this ‘hotspot’ pattern. Demonstrating its potential, MDF discovered both known and novel *trans*-eQTL in masked mouse expression data(Hasin-Brumshtein *et al*., 2016) and found known maize *trans*-regulatory relationships in a large diversity panel. Further showcasing MDF’s utility to map *trans*-regulatory relationships, we combined MDF with our F1 parent-offspring differentially *trans*-regulated ACRs to study heterotic gene expression. As parental haplotypes, and *cis*-regulatory states, are additively inherited in F1s, F1 non-additive phenotypes are expected to result from inter-haplotype *trans*-regulation. Therefore, studying these *trans*-regulatory variants, provides a good avenue to explore the underpinnings of maize seedling heterosis.

In seven-day-old maize seedlings, growth is driven by seed reserve redistribution. MDF revealed BxO *trans*-regulatory variants near critical seedling growth genes (Table 1), revealing both the growth pathways, and specific gene variants, that associate with elevated BxO seedling vigour. These genes included nitrate responsive *cytokinin response regulator2* (Sakakibara *et al*., 1998) *PEPCK3*, whose expression patterns are consistent with a gluconeogenesis role(Shenton *et al*., 2006), and *nitrate reductase1*, the rate-limiting enzyme in nitrogen assimilation(Campbell, 1999). In aggregate, these genes suggest strong seedling growth is driven by elevated cell-division and anabolic primary metabolism, with the role of cell division in maize seedling heterosis being long appreciated(Kiesselbach, 1922; East, 1936; Guo *et al*., 2010). By highlighting variants near known anabolic pathways, our data support early maize seedling vigour being carbon ‘sink’ limited, with elevated shoot carbon utilization underpinning seedling vigour. As such, we suspect manipulating the MDF highlighted genes may increase sink vegetative strength and seedling growth. Demonstrating the promise of sink strength manipulation, alteration of ear T6P-mediated sugar utilization bolsters field-grown maize yields(Nuccio *et al*., 2015). Moreover, seedling sink limitation is consistent with less conservative crop growth being beneficial – rapid seed store consumption fuels growth, essentially betting on future energetic returns, expediting light, water and minerals acquisition, while the resiliency trade-offs are managed through farmer practices. Study of other heterotic F1 seedlings is needed to see to what degree these mechanisms are conserved – in other heterotic groups and beyond.

Beyond highlighting genes in established growth pathways, the MDF approach found *trans*-regulatory genes with unknown physiological roles. Many of these are annotated kinases or TFs, including a AT-hook and SPL TF, whose cognate motifs were overrepresented in the BxO differentially *trans*-regulated ACRs. Again, the conservation of these *trans*-regulatory variants in other stiff stalkxnon-stiff stalk F1s, or other heterotic organs, remain unknown. Nonetheless, these uncharacterized *trans*-regulatory loci make great targets for future molecular genetic study, notably CRE editing to tune expression(Rodríguez-Leal *et al*., 2017; Liu *et al*., 2021; Zhou *et al*., 2025), as they likely have large genomic and organismal growth impacts. This potential for gene discovery is the allure of MDF empowered *trans*-regulatory variant detection – by establishing *trans*-regulatory relationships, MDF can facilitate gene discovery from extant and future GWAS data, providing hypotheses to be tested in targeted studies.

MDF is species and genomic modality agnostic, allowing application to diverse biological questions, and has several advantages beyond its discovery power. Although the individual *trans*-relationships supporting each SNP are still prone to false positives, MDF uncovers many such associations, providing information about the variant’s biological function. Also, MDF exploits genetic associations, potentially uncovering spatial/temporal disparate relationships. We anticipate MDF’s increased discovery power to be especially useful in emerging single-cell QTL studies(Benaglio *et al*., 2023; Marand *et al*., 2025), which suffer from a further exacerbated testing burden. MDF establishes *trans*-regulatory relationships, enabling better understanding of how gene regulatory networks elicit phenotypes and providing gene candidates for 21^st^ century trait manipulation.

## Methods

### Plant growth, nuclei isolation, Tn5 tagmentation, library prep and sequencing

Hybridization hand crosses were conducted on glasshouse grown plants in 2023 from seed received from the maize CO-OP stock center. Each F1 genotype analysed was the same hand cross. All analyzed plants were grown in the same conditions and space as those in Marand et al., (2025). Specifically, all biological replicates (n=4-6) were grown in Sungro soil (Sungro Horticulture Canada Ltd) under a 16-hour photoperiod (50:50 mixture of fluorescent light from 4100K Sylvania Supersaver Cool White Delux F34CWX/SS, 34W and 3000K GE Ecolux w/ starcoat, F40CX30ECO, 40W lighting), at a constant ∼25°C. All genotypes were grown concurrently, in trays that were blocked on genotype. Seven days after planting, the first node +/-∼1cm of tissue was harvested for fresh nuclei extraction with a Personna razor (74-0002 Personna Stainless Steel Surgical Prep Blades) between 8 and 10am (1-3 hours post dawn). Nuclei extraction was done as before(Marand *et al*., 2025). Briefly, the isolated tissue was chopped over ice for 2 minutes with Personna razor blades in nuclei extraction buffer (1X NIB: 10mM MES:KOH pH 5.4, 10mM NaCl, 10mM KCl, 2.5 mM EDTA pH 8, 250 mM sucrose, 0.1mM spermine, 0.5mM spermidine, 1mM fresh DTT) with 0.5% vol/vol Triton X100. Then this tissue slurry was passed though 40 μm cell strainer (Pluriselect) into a 2mL conical tube and spun at 4°C and 500g for 5 minutes in a swinging bucket centrifuge (Eppendorf 5810R Centrifuge) to pellet the nuclei. The supernatant was then removed, and the nuclei resuspended in 500 μL 1X NIB before being passed through a 10 μm cell strainer (Pluriselect) before being gently layered on top of 1mL of 35% vol/vol Percoll:1XNIB in another 2mL conical tube and spun for 10 minutes at 4°C and 500g. Afterwards, all supernatant was removed, and the nuclei were resuspended in 1X TAPS buffer (10mM TAPS-NaOH, pH8.0, 5mM MgCl2), stained with DAPI (1μg/mL final concentration) before being counted. Per replicate, ∼40 thousand nuclei were then Tn5 tagmented in 2.5X TAPS as previously described, except using 1.75μg Tn5(Ricci et al., 2019). These tagmentation products were then purified (Monarch PCR & DNA Cleanup Kit; as dsDNA <2kb; NEB #T1030), and PCR amplified, as described previously (Ricci et al., 2019), to add sequencing adapters. These libraries were the cleaned using AMPure XP beads (Beckman Coulter) before sequencing (∼100 million 150 bp paired end reads) on an Illumina Novaseq 6000.

### ATAC-seq read processing

FASTQC was used to confirm the quality of all sequences libraries. Then, all paired end reads were aligned to either a single (B73) or concatenated reference that corresponded to the genotypes in the parent-offspring comparison(Hufford *et al*., 2021). These concatenated references are the result of merging the reference chromosome sequences (unplaced contigs excluded) and annotations, after adding a unique identifier to the chromosome fields of each haplotype (chr1 –> B73_chr1 and Oh43_chr1). For the single reference, reads were trimmed with Trimmomatic (Bolger *et al*., 2014), aligned using bowtie2(Langmead and Salzberg, 2013) (--no-unal), filtered for unique alignments, and deduplicated via Picardtools and then filtered against a blacklist. Tn5 fragments and insertion sites were then quantified via Sinto and a dummy cell barcode that was otherwise unused. Peaks (ACRs) for the single reference were those previously generated, and annotated, from the single-cell ATAC-seq data in Marand et al. (2025).

Alignment to the concatenated references has been described elsewhere(Engelhorn *et al*., 2025). After trimming via Trimmomatic, the reads are aligned via the spliced read aligner STAR(Dobin *et al*., 2013) (--outSAMmultNmax 2 –-winAnchorMultimapNmax 100 –-alignIntronMax 1 –-outBAMsortingBinsN 5), to account for indels between the references. Then these alignments are treated as the single reference reads to generate Tn5 insertions. From these insertions, MACS2(Zhang *et al*., 2008) was used to call peaks (g 1.6e9 –-nomodel –-keep-dup all –-extsize 150 –-shift –50 –-qvalue 0.05). These MACS2 peaks were then cleaned using a perl script (cleanBED.pl). To generate reproducible peaks for each concatenated reference, we first merged all overlapping peaks found in all F1 replicates. We then retained these merged peaks for analysis if there was an overlapping peak in n-1 of the F1 biological replicates.

### ACR Homology/conservation, inheritance, positional and motif content analysis

After defining a peak (ACR) set for each parent-offspring pair, we used a haplotype reciprocal alignment(Li *et al*., 2009; Quinlan and Hall, 2010; Neph *et al*., 2012; Li, 2018) approach to determine intra-maize ACR conservation. Minimap2(Li, 2018) (-x splice –t 20 –k 12 –a –p 0.4 –N 3) was used to align each ACR to the other haplotype reference (e.g. a B73 ACR aligned to the Oh43 genome). Then, exploiting the pan-gene annotation(Hufford *et al*., 2021), we filtered for alignments that shared at least one closest gene to the query ACR (e.g. the Oh43 alignments that were closest to the same pangene as the query B73 ACR). We then intersected the alignment coordinates with the ACR complement found on the other haplotype (e.g. comparing the B73-to-Oh43 alignment coordinates to the ACRs found on the Oh43 haplotype), requiring an alignment that spanned at least 45% of the ACR detected on the other haplotype. If this 45% sequence identity relationship was reciprocal (e.g. the B73-to-Oh43 alignment finds a given Oh43 ACR, and using that given Oh43 ACR in a Oh43-to-B73 alignment finds our original B73 ACR), we classified those ACRs as ‘shared’. However, if the 45% sequence identity threshold was not reciprocal, these ACRs were classified as ‘ambiguous.’ ACRs with no 45% sequence identity alignment next to the same gene in the other reference were classified as ‘unique.’ Finally, ACRs with more than one 45% identity alignment on the other haplotype, next to at least one of the same genes, were classified as ‘duplicated’ ACRs.

To classify ACR chromatin accessibility inheritance patterns, we mirrored the gene expression inheritance classification in Coolon et al., (2014). However, we modified this approach to be more stringent about calling ACR Tn5 insertions rates ‘different’ between genotypes. Specifically, similar ACRs had were either alike in chromatin accessibility between all genotype comparisons (e.g. B73 to BxK, Ki3 to BxK, B73 to Ki3; DESeq2(Love *et al*., 2014); minimum p>0.05) or had a fold change of less than two in each parent-to-offspring comparison. For the rest of the ACRs, we used the differences in chromatin accessibility between parent and F1 offspring to classify ACR inheritance: Dominant ACRs had fold change <1.25 and >1.25 relative to each parent, additive ACRs had increased chromatin accessibility relative to one parent, but lower than the other, under-dominant ACRs had lower chromatin accessibility than both parents and over-dominant ACRs higher chromatin accessibility than both parents.

ACRs were also categorized based upon their position relative to genes as previously described(Ricci *et al*., 2019; Marand *et al*., 2025). Using the corresponding gene annotation for each reference genome, ACRs overlapping all annotated TSS (+/-25bp on either strand) were categorized as ‘TSS,’ those farther than 25bp away, but less than 2kb were categorized as ‘proximal,’ those overlapping annotated genes, but not the TSS, were categorized as ‘genic’ and those >2kb away from genes ‘intergenic.’ To find relative enrichment/depletion of these ACR categories to one another, we compared the proportion of ACR category intersection (e.g. shared ∩ genic) to the proportion of the ACR categories in the entire ACR set (outlined in Figure 2).

All TF F1 ACR motif analysis was conducted on the concatenated reference ACR sets, as these most accurately reflect the CREs available for TF binding. First, for each parent-offspring comparison, the occurrence of all motifs listed in the JASPAR 2024 database(Rauluseviciute *et al*., 2024) was found via FIMO(Grant *et al*., 2011) (--max-stored-scores 500000). Then, the rate motif occurrence was compared between that found in an ACR category, to that expected due to chance (outlined in Figure 3). For example, the observed proportion of the TCP23 motif (MA1066.1) in additive ACRs was compared to the proportion of all motifs found in additive ACRs. The difference between these proportions (observed – expected) is the measure of effect size that was used for visualization. Motif enrichment was also conducted on the ACRs that trans-associated with the two MDF-passing variants within the *Tb1* control region. *De novo* motif enrichment was conducted using STREME(Bailey, 2021). Motif enrichment was done via SEA(Bailey and Grant, 2021), using TB1 motif MA1430.2.

### Narrow sense heritability estimates

Narrow sense heritability (h^2^) estimates were conducted as outlined elsewhere(De Villemereuil *et al*., 2013). Briefly, for each B73-reference ACR defined in Marand et al. (2025), chromatin accessibility was first compared via DESeq2(Love *et al*., 2014) between all sampled genotypes and those with at least one significant difference (FDR > 0.05) were retained for h^2^ assessment. This excludes ACRs with no genetic or environmental variation in our sample. To ensure adequate Tn5 insertions (i.e. to limit technical bias due to sequencing depth variation), ACRs were then filtered to retain those with >0.1 CPM Tn5 insertions. After these filters, the observed F1 chromatin accessibility value was compared to that predicted based on the mid-parental value. This was done by correlating the F1 biological replicate values with the mid-parental value. ACRs with a negative correlation were set to an h^2^ of zero(De Villemereuil *et al*., 2013). ACRs were then split into three h^2^ groups: h^2^<0.3 low heritability, h^2^ 0.3-0.65 ‘transcriptional’ heritability, h^2^>0.65 ‘regulatory’ heritability. Gene ontology enrichments were then conducted on the genes closest to these grouped ACRs through the ‘TopGO’ R package (Alexa and Rahnenfuhrer, 2010).

### Detection of F1 differentially *trans*-regulation over ACRs

As outlined in the main text, this analysis looked for allele-specific chromatin accessibility that deviated from the ‘half-parental’ value consistent with allelic inheritance and strictly *cis* control of chromatin accessibility. For unique ACRs, we first filtered out ACRs called unique due to potential false negative peak calling in one genotype. To do this, unique ACRs with >0.15 CPM Tn5 insertions in the peak-less inbred samples were removed. For example, in the BxK comparison, a B73 unique ACR would be excluded if the Ki3 inbred sampled had >0.15 CPM Tn5 insertions cross-mapping to the ACR on the B73 haplotype. This >0.15 CPM Tn5 cut-off was determined empirically based on the aggregate of all unique ACRs in all F1 concatenated genomes. As elsewhere, unique ACRs were then filtered to retain those with >0.1 CPM Tn5 insertions to minimize the technical effects of differing read depth.

For the retained unique ACRs, all Tn5 insertion CPMs were log transformed and the observed F1 value compared to the halved parental value. To detect *trans*-regulated ACRs at a range of CPM, this deviation or ‘distance’ from the ‘half parental’ value was normalized based on F1 CPM of the ACR. Then, this normalized distance was plotted for all ACRs across all F1 groups. Based on this aggregate normalized distance, a stringent cut off was set: a ACR was deemed F1 differentially *trans*-regulated if its normalized distance was greater than two standard deviations away from the whole population mean in n-1 F1 biological replicates and the F1 mean.

A similar approach was used for Tn5 fragment reads over SNPs in shared ACRs. First, reciprocal minimap2(Li, 2018) alignment was conducted with similar parameters (-x splice –-secondary=no –t 20 –k 12 –a –-MD –p 0.4 –N 1) as those used to initially identify the shared ACRs, but just using the two reference sequences from the two shared ACRs (e.g. BxK shared ACR would align the corresponding ACRs from the B73 and Ki3 reference). From this alignment, a python script extracted all SNPs and their ACR positions into a ‘key’ file. This key file when then used to find SNPs called in the same position, and with the cognate reference/alternative base call, for the two reciprocal ACR alignments. Then, per shared SNP, the number of reads containing the appropriate SNP were counted from the corresponding alignment files using lightly modified JVarkit (Lindenbaum and Redon, 2018) scripts and CPM normalized.

After generating SNP-resolved Tn5 fragment counts, we used the same approach as outlined for unique ACRs. Specifically, we again measured the deviation or ‘distance’ from the ‘half parental’ value, which was again normalized for CPM. These normalized distances were aggregated across all F1 groups, and a *trans*-regulated cut-off was set based off values greater than two standard deviations from the aggregate population mean. Finally, SNPs were considered F1 differentially *trans*-regulated if their normalized distance was greater than this cut-off, both in n-1 F1 biological replicates and the F1 mean.

### *Trans* chromatin accessibility genetic association

*Trans* chromatin accessibility QTL and gene expression QTL were detected with tensorQTL v1.0.09(Taylor-Weiner *et al*., 2019) with non-default settings (--dosages –-pval_threshold 0.01 –-mode trans) using the same input files (covariates, and bulk tissue normalized chromatin accessibility values) as previously reported for cis-caQTL detection(Marand *et al*., 2025). To reduce the number of tests with independent filtering, we only tested SNPs that additionally colocalized with a non-genic ACR, resulting in a total of 481,944 variants. The same variant set was used for both trans caQTL and eQTL discovery.

### Multiple discovery filtering

MDF revolves around the notion that unlinked (>5Mpb away) *trans*-association false positives are independent events. This independence allows for the determination of the probability of observing n, or more, false positive variant-by-ACR/gene associations at a given α, per variant tested (Equation #1).

Multiplying this per-SNP-false-positive-probability by the total number of variants tested provides the expected false-positive number of variants that associate a repeat discovery number (n-DISC) of ACRs at an α threshold, assuming genotype and environment remain unconfounded(Kaur *et al*., 2024). After filtering the observed data against these n-DISC and α thresholds, we calculate a false positive rate (predicted false positives / observed positives). MDF passing variants have more trans associations than that expected due to chance alone, suggesting they are linked to true positive *trans*-regulatory genes.

## Acknowledgements

This research was supported by the National Science Foundation (IOS-2026554 and IOS-1856627) to RJS as well as the UGA Office of Research to RJS. A.P.M. was supported by the National Institute of General Medical Sciences of the National Institutes of Health (1R00GM144742). We would like to thank Micheal Boyd and Noah Herrington for growth facility maintenance and unparalleled plant care, as well as the staff at the Georgia Advanced Computing Resource Center (GACRC) for their tireless support of our computing cluster. We also thank our talented lab technicians Yinxin Dong and Yangyang Xu for their outstanding efforts that keep the lab running smoothly.

